# Tensor decomposition reveals coordinated multicellular patterns of transcriptional variation that distinguish and stratify disease individuals

**DOI:** 10.1101/2022.02.16.480703

**Authors:** Jonathan Mitchel, M. Grace Gordon, Richard K. Perez, Evan Biederstedt, Raymund Bueno, Chun Jimmie Ye, Peter V. Kharchenko

## Abstract

Tissue- and organism-level biological processes often involve coordinated action of multiple distinct cell types. Current computational methods for the analysis of single-cell RNA-sequencing (scRNA-seq) data, however, are not designed to capture co-variation of cell states across samples, in part due to the low number of biological samples in most scRNA-seq datasets. Recent advances in sample multiplexing have enabled population-scale scRNA-seq measurements of tens to hundreds of samples. To take advantage of such datasets, here we introduce a computational approach called single-cell Interpretable Tensor Decomposition (scITD). This method extracts “multicellular gene expression patterns” that capture how sample-specific expression states of a cell type are correlated with the expression states of other cell types. Such multicellular patterns can reveal molecular mechanisms underlying coordinated changes of different cell types within the tissue, and can be used to stratify individuals in a clinically-relevant and reproducible manner. We first validated the performance of scITD using *in vitro* experimental data and simulations. We then applied scITD to scRNA-seq data on peripheral blood mononuclear cells (PBMCs) from 115 patients with systemic lupus erythematosus and 56 healthy controls. We recapitulated a well-established pan-cell-type signature of interferon-signaling that was associated with the presence of anti-dsDNA autoantibodies and a disease activity index. We further identified a novel multicellular pattern linked to nephritis, which was characterized by an expansion of activated memory B cells along with helper T cell activation. Our approach also sheds light on ligand-receptor interactions potentially mediating these multicellular patterns. As validation, we demonstrated that these expression patterns also stratified donors from a pediatric SLE dataset by the same phenotypic attributes. Lastly, we found the interferon multicellular pattern and others to be conserved in a COVID-19 dataset, pointing to the presence of both general and disease-specific patterns of inter-individual immune variation. Overall, scITD is a flexible method for exploring co-variation of cell states in multi-sample single-cell datasets, which can yield new insights into complex non-cell-autonomous dependencies that define and stratify disease.

## Introduction

Gene expression is a defining feature that distinguishes different cell types. However, gene expression profiles derived from the same cell type can also vary across individuals, driven by a combination of genetics and environment. Most often, gene expression is compared between individuals in case-control studies or is used to infer sample subgroupings. Alternatively, studies such as the Genotype-Tissue Expression (GTEx) project have revealed the genetic basis of tissue-specific gene expression by mapping natural genetic variation associated with expression differences in human populations.^1,2^ Applications of single-cell RNA-sequencing (scRNA-seq) have so far focused on the characterization of transcriptional differences among different cell types and cell states, and analysis of inter-individual variation has been hampered by small sample sizes and the presence of technical batch effects that are often difficult to separate from biological variation. Experimentally, this stimulated the development of multiplexed designs where samples from multiple individuals could be profiled in one run, thereby reducing confounding by technical batches.^3,4^ Analytically, a variety of approaches have been developed to perform dataset alignment in order to overcome technical variation from non-multiplexed datasets.^5–8^ These tools establish correspondence between samples, effectively treating both technical and biological differences between individuals as a problem to be overcome. However, biological differences (e.g., between cases and controls or genetically different individuals) are often central for the downstream interpretation, but relatively few approaches exist to explore them systematically.

The existing methods to compare single-cell transcriptomes between individuals generally require pre-defined groups of samples such as in case/control studies. Comparisons between sample groups can be made using cell-level data or aggregated counts for all cells within a given type or cluster (often referred to as a “pseudobulk” operation).^9,10^ After computing the pseudobulk profiles, differential expression (DE) tools designed for bulk RNA-seq analysis can be used to compare sample groups one cell type/cluster at a time. Pseudobulk analysis has been demonstrated to provide a superior balance of robustness, performance, and runtime^10,11^ and has been used in several single-cell case-control studies to date.^3,12–16^

However, case-control DE approaches are unable to stratify patients into subgroups and do not account for expression dysregulation that may occur jointly in multiple cell types. Patient subgrouping is of interest when the covariate defining a group has not been well-captured (e.g., disease status has been only partially recorded or recorded with errors) or when additional heterogeneity exists across samples. Standard matrix decomposition methods such as principal components analysis (PCA) and non-negative matrix factorization can, in principle, be used to describe gene expression variation across samples one cell type/cluster at a time. However, as we will illustrate, it is more informative to consider inter-individual variation within multiple cell populations jointly. The structure of scRNA-seq data uniquely enables such analysis since it yields gene expression information for multiple cell types per sample. Such a joint decomposition would reveal subsets of samples that exhibit gene expression dysregulation in multiple cell types. For example, samples undergoing an innate immune response would display increased macrophage chemoattractant secretion and neutrophil migration, as these processes are coordinated. These types of multicellular patterns may reflect cell-type-specific responses to the same external signals and the complex interactions between cells. As a result, this type of approach will naturally help us identify sample-specific cell-cell interactions and allow us to generate informed hypotheses regarding their potential downstream effects. Overall, by extracting such patterns of gene expression variation, we hypothesize that we can better characterize the molecular bases of complex phenotypes.

Here, we developed an unsupervised method for analysis of inter-individual variation, called single-cell Interpretable Tensor Decomposition (scITD). The method infers multicellular patterns of gene expression in scRNA-seq datasets containing many patients or samples (Figure 1A). We define a “multicellular pattern” to be a collection of genes in various cell types that co-vary together across samples. These multicellular patterns can be further associated with known covariates (e.g., disease, treatment, or technical batch effects) to reveal clinically relevant stratification of heterogeneous patient cohorts. We first assessed the performance of scITD using simulated and real data from an *in vitro* experiment. Then, we applied scITD to investigate inter-individual heterogeneity in peripheral blood mononuclear cell (PBMC) expression using a dataset with 115 systemic lupus erythematosus (SLE) patients and 56 healthy controls. SLE is a heterogeneous autoimmune disease that can manifest with a wide array of symptoms and has few available targeted therapies ^17,18^. We identified six multicellular patterns of gene expression that stratify SLE patients, and we show that these are associated with clinical variables including disease activity and nephritis, one of the most severe complications of SLE. These patterns were examined in depth to identify channels of intercellular communication and changes in cell-type composition and were validated using a pediatric SLE dataset. We further compared multicellular patterns derived from adult and pediatric SLE datasets as well as those from a COVID-19 dataset, revealing both conserved- and disease-specific immune responses.

**Figure 1.**
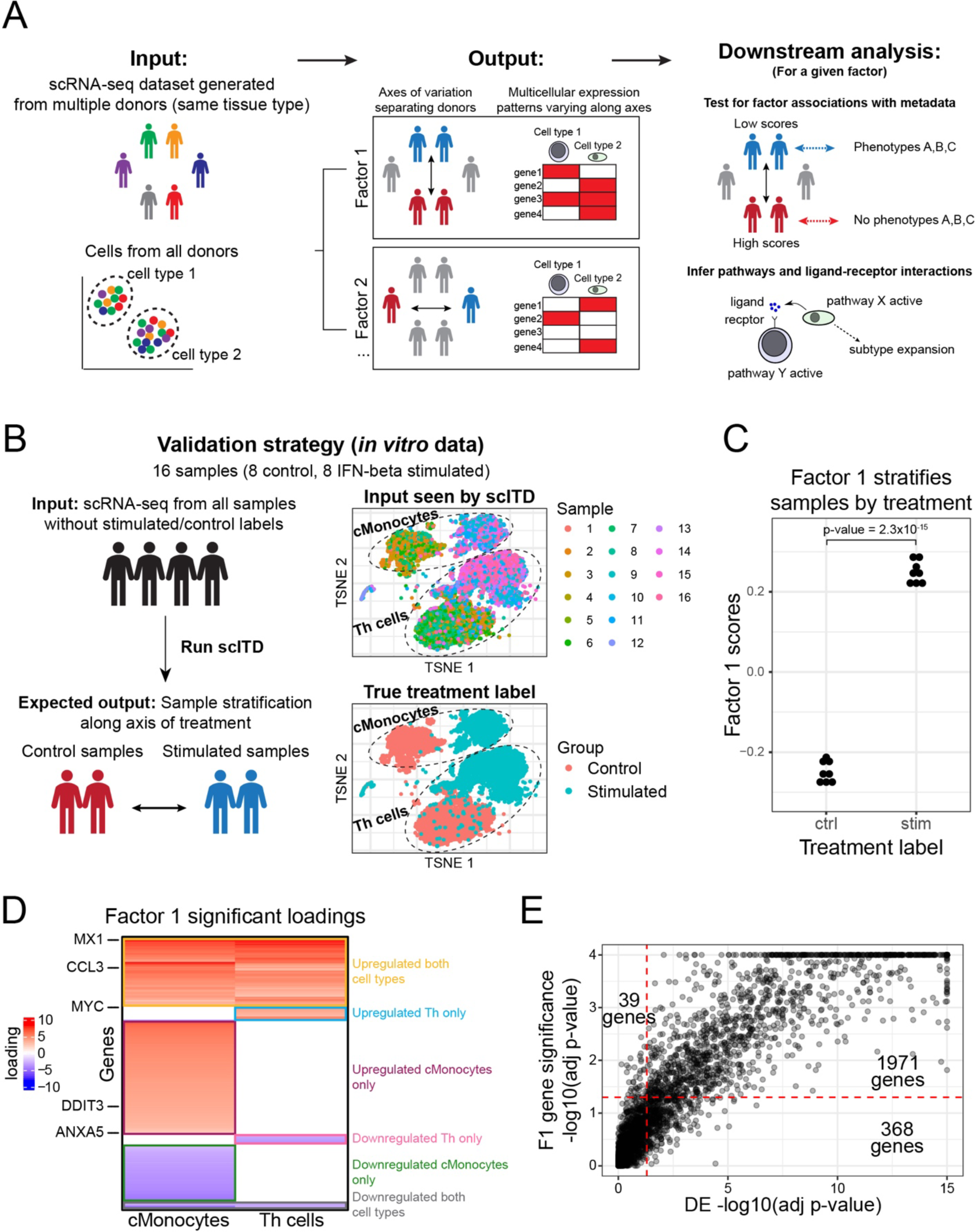
General overview of scITD and demonstration of functionality. (A) The overall goal of scITD. The tool takes as input a scRNA-seq UMI count matrix as well as a metadata matrix with cell type and donor annotations (left). scITD then identifies axes of expression variation using data from multiple cell types jointly to stratify donors (middle). These axes and patterns can be further analyzed for metadata associations and for signatures of biological processes (right). (B) Validation strategy using data from IFN-beta stimulation experiment (left). Also shown are t-SNE plots of the non-sample-aligned data colored by sample or by condition (right). (C) The sample scores for factor 1 after applying scITD to the IFN-beta stimulation dataset. Samples are grouped by their true treatment labels. A linear model F test was used to compute significance of the association. (D) Loadings heatmap for factor 1. Only loadings for genes that are significantly associated with factor 1 scores in either cell type are shown (Methods). Genes only significant in one cell type are given the color white in its non-significant cell type. Rows of the loadings heatmap are hierarchically clustered. Boxes indicate groups of genes found to be up/down regulated in a combination of cell types. Up/down direction labels are for treated samples relative to the control samples. (E) Comparison of DE adjusted p-values (from S1B) to gene expression-factor 1 sample score association p-values (Methods). Each point is a gene in one of the two cell types. The dashed red lines are located at an adjusted p-value of 0.05.

## Results

### Approach and evaluation of performance

To extract multicellular patterns that vary across individuals, we first generate normalized pseudobulk expression profiles per sample per cell type (Methods). When *C* cell populations from *N* samples are collapsed into pseudobulk profiles, the dataset can be represented as a 3-dimensional matrix – a tensor *T* with dimensions *N* × *G* × *C*, where *G* is the number of genes (Figure S1A left). As we intend to capture sample expression differences within cell types as opposed to cell type markers, we scale and center the expression of each gene (in each cell type) across individuals. We then apply the Tucker tensor decomposition^19^ to extract the *K* most informative factors, which represent axes of variation stratifying individuals. Each factor consists of a gene-by-cell type matrix of loadings values, representing a multicellular gene expression pattern as well as a vector of sample scores, indicating the relative degree to which the respective pattern is expressed in each sample (Figure 1A middle and S1A). Specifically, the sample scores vector for a given factor is analogous to an eigengene for gene co-expression modules,^20^ as it describes each sample’s weighted average expression of genes in the corresponding loadings matrix. Therefore, we can use them as any other continuous variable to test them for correlations with other patient metadata (Figure 1A right).

Before testing scITD on a dataset from a heterogeneous patient cohort, we aimed to demonstrate that it could accurately stratify samples with a single known source of expression variation across samples. For this analysis, we chose a dataset of 16 samples of PBMCs from SLE patients, half of which were stimulated *in vitro* with interferon-beta (IFN-beta)^3^ (Figure 1B). The goal of this analysis was to provide scITD the data without treatment labels and to verify that the algorithm could separate samples along the treatment axis. Additionally, we aimed to show that the multicellular pattern stratifying samples in this manner would recapitulate the expression differences that would be obtained by traditional differential expression tests between treatment and control groups for any given cell type. For simplicity of the demonstration, we limited the analysis to just classical monocytes (cMonocytes) and CD4+ T cells (Th cells). In this analysis we conservatively assume that the 16 samples are independent even though in reality each donor had one treated and one control sample. After applying scITD to this dataset, we found that the first factor perfectly stratified the samples by treatment (Figure 1C). The corresponding gene loadings matrix reveals a pattern consisting of both shared and cell-type-specific genes responsive to IFN-beta stimulation (Figure 1D). To interpret these results in terms of attributing expression differences to samples, one should consider the signs of the sample scores as well as those of the gene loadings. For a given factor, samples with positive scores have higher expression of positive loading genes compared to samples with negative scores, and vice versa. Out of the 2507 genes tested (overdispersed genes included in the tensor), we found 1364 genes to be significant in the monocytes and 653 genes to be significant in the CD4+ T cells (Padj<0.05). A direct differential expression (DE) test between stimulated and control samples revealed a highly similar pattern with many of the same cell-type-specific effects on expression (Figure S1B). Out of the 2017 significant gene cell type pairs that were also tested in the DE analysis, we observed that 98% (1971/2010) were significant in both analyses (Figure 1E). The Benjamini-Hochberg (BH) procedure was used for multiple hypothesis test correction here as well as for other p-value adjustments throughout the study.^21^ Similarly, we observed a high correlation between the factor loadings for all genes and the log fold-change values from the DE analysis and demonstrated that this correlation remains robust after downsampling the data to varying degrees (Figure S1C and S1D). This example demonstrates that scITD can accurately stratify samples when the underlying gene expression differences involve both shared and cell-type-specific genes.

To further evaluate the performance of scITD, we simulated a scRNA-seq dataset with 40 donors (1 sample per donor) and two cell types (Figure S1E left). The dataset was designed to include two multicellular patterns that varied across donors (Figure S1E right), and each pattern involved mostly different genes in each cell type. scITD accurately prioritized the relevant donors and genes involved in each pattern (Figure S1F). We also developed several approaches to help determine the appropriate number of factors into which the initial tensor should be decomposed including a significance test based on SVD (Methods). When applying these approaches to the simulated dataset, they recommended the dataset be decomposed into two factors, accurately reflecting the number of ground truth multicellular patterns encoded into the dataset (Figure S1G, S1H, and S1I).

### Analysis of an SLE dataset identifies novel multicellular patterns that stratify patients

Next, we prepared a large scRNA-seq dataset^22^ of PBMCs from 115 SLE patients and 56 healthy donors for analysis with scITD (Figure 2A). Our primary goal was to identify multicellular patterns that stratified patients to help explain the vast patient-to-patient phenotypic heterogeneity seen in this disease. Therefore, we initially limited our analysis to patients only. We also focused our analysis on 7 of the major cell types annotated at a coarse-grained level (Figure 2B left). After transforming expression counts to pseudobulked counts, we further applied batch correction, as groups of donors were pooled and processed together in different 10X Chromium lanes (Methods). Before running scITD, we needed to determine an appropriate number of factors to decompose the dataset into. We applied our SVD-based rank determination approach and others (Methods), which suggested we use somewhere between 7-12 factors (Figure S2A and S2B). Conservatively, we applied scITD to extract only 7 factors. Supporting our parameter choice, we noted that these factors were highly stable even when randomly downsampling donors from the start of the pipeline (Figure S2C).

**Figure 2.**
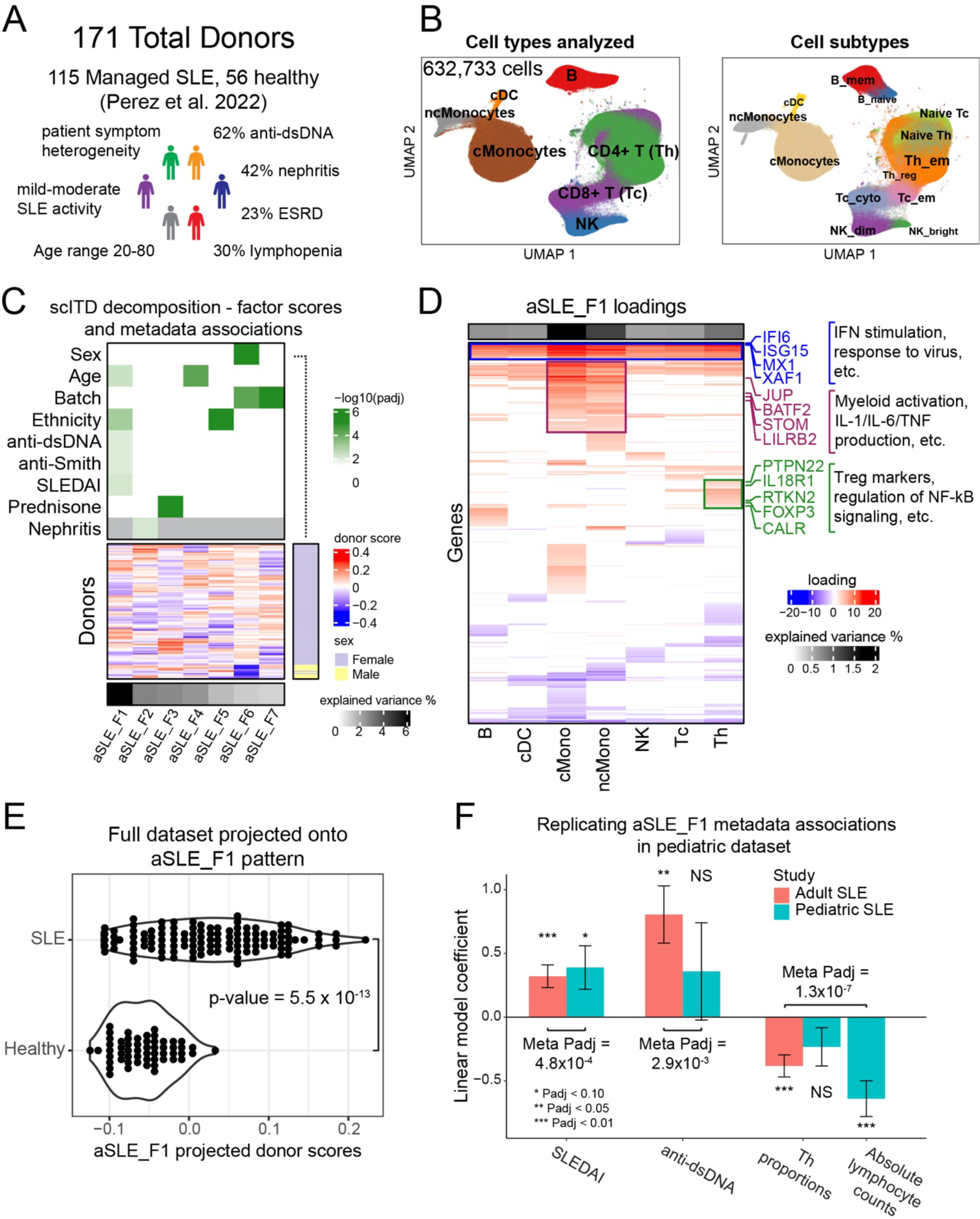
SLE scRNA-seq dataset overview and scITD analysis. (A) Description of the SLE PBMC dataset with summary statistics for select clinical attributes. (B) UMAP of single-cell gene expression from the SLE dataset, showing the coarse clustering used in the downstream analyses (left) and cell subtype annotations (right). (C) Donor scores heatmap with metadata association p-values annotated at the top. The p-values were calculated using univariate linear model F-tests. Rows are hierarchically clustered. Columns are ordered by explained variance for each factor, and this is displayed at the bottom of the heatmap. (D) Loadings matrix for factor aSLE_F1 limited to only significant genes. The top annotation shows the percent of overall explained variance for each cell type of the factor. The genes highlighted with different colors are a few leading-edge genes for the gene sets with corresponding colors. Rows are hierarchically clustered. (E) The full SLE dataset (including healthy donors) projected onto aSLE_F1, and the association between these projected scores and SLE status. The p-value was calculated using a linear model F-test. (F) Replicating aSLE_F1 metadata associations in pediatric SLE dataset. The pediatric dataset was projected onto the aSLE_F1 multicellular pattern to yield projected scores, which were used for metadata association tests with this dataset. Univariate linear model F-tests were used to evaluate statistical significance. The Fisher method was used for meta-analysis of aSLE_F1 associations with cognate phenotypes across datasets indicated by the brackets.

We observed several significant associations between the factor donor scores and metadata variables such as sex, ethnicity, age, and disease activity (SLEDAI score) among others (Figure 2C). For this dataset, factors are labeled as aSLE_F1, aSLE_F2, etc. to indicate they are derived from the adult SLE (aSLE) dataset. We first investigated aSLE_F1, which had associations with SLEDAI score and autoantibody presence (Figure 2C). The loadings matrix for this factor (Figure 2D) revealed that donors with high aSLE_F1 scores had increased expression of interferon-stimulated genes (ISGs) in multiple cell types. Directionally, these individuals also tended to have higher SLEDAI scores and autoantibody presence (Figure 2F). The connection between higher ISG expression and these clinical features has been previously shown in several studies, and ISG is reportedly overexpressed in roughly half of all SLE patients.^23–30^ Specifically, the Pearson correlation between aSLE_F1 donor scores and SLEDAI was 0.321, which is similar to what has been reported in other studies.^31–33^ The aSLE_F1 factor also consisted of myeloid-specific activation genes and Th-specific genes, which likely represent cell-type-specific responses to interferon stimulation and other co-occurring processes (Figure 2D). When projecting the full dataset (including healthy donors) onto this multicellular pattern, it clearly separated out the healthy donors from the subset of SLE patients with high ISG expression (Methods) (Figure 2E). This observation aligned with our expectation for healthy individuals, as ISGs are normally upregulated during infections. Further, ISG expression is not a perfect disease biomarker as a subset of SLE patients had low ISG expression comparable to the healthy donors. We further applied gene set enrichment analysis (GSEA) per cell type for this factor (Figure S2D), and this yielded the expected enrichment of the “response to type I interferon” gene set in all cell types as well as other gene sets enriched in specific cell types, especially monocytes. Some of the monocyte-specific gene sets included interleukin production and TNF production among others. Interestingly, the Th cells for the aSLE_F1-high SLE patients also had upregulation of the canonical Treg marker FOXP3 (Figure 2D). Consistent with this, we observed a significant increase in Treg cell proportions for donors aSLE_F1-high donors (Figure S2F). Previous studies have also shown increased numbers of Tregs in SLE patients compared to healthy donors and often accompanying high ISG expression.^34,35^ In addition to this finding, we also found a significant reduction in the proportion of total Th cells for aSLE_F1 high donors (Figure S2E).

Next, we set out to validate and compare these results with a pediatric SLE scRNA-seq dataset. This secondary dataset consisted of PBMC scRNA-seq data from 33 pediatric SLE cases and 11 pediatric healthy controls (Figure S2G).^30^ We first aimed to test whether our aSLE factors would also stratify the pediatric SLE (pSLE) patients by the same metadata associations. Therefore, we projected the pediatric data onto the aSLE multicellular patterns to get projected scores for the pediatric donors. We then tested the projected scores for associations with the same metadata. For aSLE_F1, we observed that all metadata associations had preserved directionality and yielded significant p-values by Fisher meta-analysis (Figure 2F). The consistency of these associations in both adult and pediatric cohorts bolsters the connection between the aSLE_F1 multicellular pattern and these metadata, in particular reduction in the proportion and abundance of T lymphocytes. While the original pediatric study identified the association between SLEDAI and ISG expression, they did not report the association between lymphocyte counts and ISG expression as we demonstrated. Further supporting the connection between elevated ISGs and T lymphopenia, treatment by Anifrulomab, a recently approved monoclonal antibody specific for subunit 1 of the type 1 interferon alpha receptor (IFNAR1) restores T lymphocyte counts.^36^ This highlights the ability of scITD to reveal new connections between gene expression patterns and patient phenotypes. In a separate analysis, we also applied scITD to stratify the pediatric patients. This yielded three factors, pSLE_F1, pSLE_F2, and pSLE_F3, that stratified patients in a highly similar manner to aSLE_F1, aSLE_F2, and aSLE_F3 from the adult SLE dataset (Figure S2H). This demonstrates that the major sources inter-individual expression variation in SLE are highly conserved across these two patient populations.

Lupus nephritis is one of the most severe complications of SLE, and anti-dsDNA autoantibodies are a critical though insufficient component to its development.^37^ Therefore, we sought to identify multicellular patterns that are associated with nephritis when autoantibodies are present. Factor aSLE_F2 exhibited a significant association with the frequency of lupus nephritis among SLE patients who were positive for anti-dsDNA autoantibodies (Figure 3A). The frequency of patients with nephritis was computed using a sliding window along the factor scores to calculate the number of patients positive for lupus nephritis among those positive for anti-dsDNA autoantibodies (Methods). Notably, there is no association in patients negative for anti-dsDNA autoantibodies, and co-occurrence of anti-smith autoantibodies with lupus nephritis for this factor was less significant (p-value ∼ 0.05). Projecting the pediatric data onto aSLE_F2 replicated the expected significant metadata associations with nephritis but also identified an additional association to creatinine levels (Figure 3C and S3A). Analysis of the loadings matrix for aSLE_F2 revealed a coordination of multiple cell-type-specific processes with a rough grouping of processes by T cells, myeloid cells, or B cells (Figure 3B). GSEA analysis and evaluation of top dysregulated genes point to processes likely perturbed in these cell types. For instance, genes such as *G0S2* were upregulated in myeloid cell types, potentially suggesting increased myeloid proliferation. Additionally, *CXCR5* was upregulated in B cells, which is possibly indicative of B cell trafficking to germinal centers. Further, Th cells expressed overexpressed genes for p38 MAPK signaling and downregulated TCR genes, indicating likely T cell activation. The observation of increased p38 MAPK signaling was particularly intriguing because there is evidence that this pathway may play a causal role in the development of lupus nephritis.^38,39^

**Figure 3.**
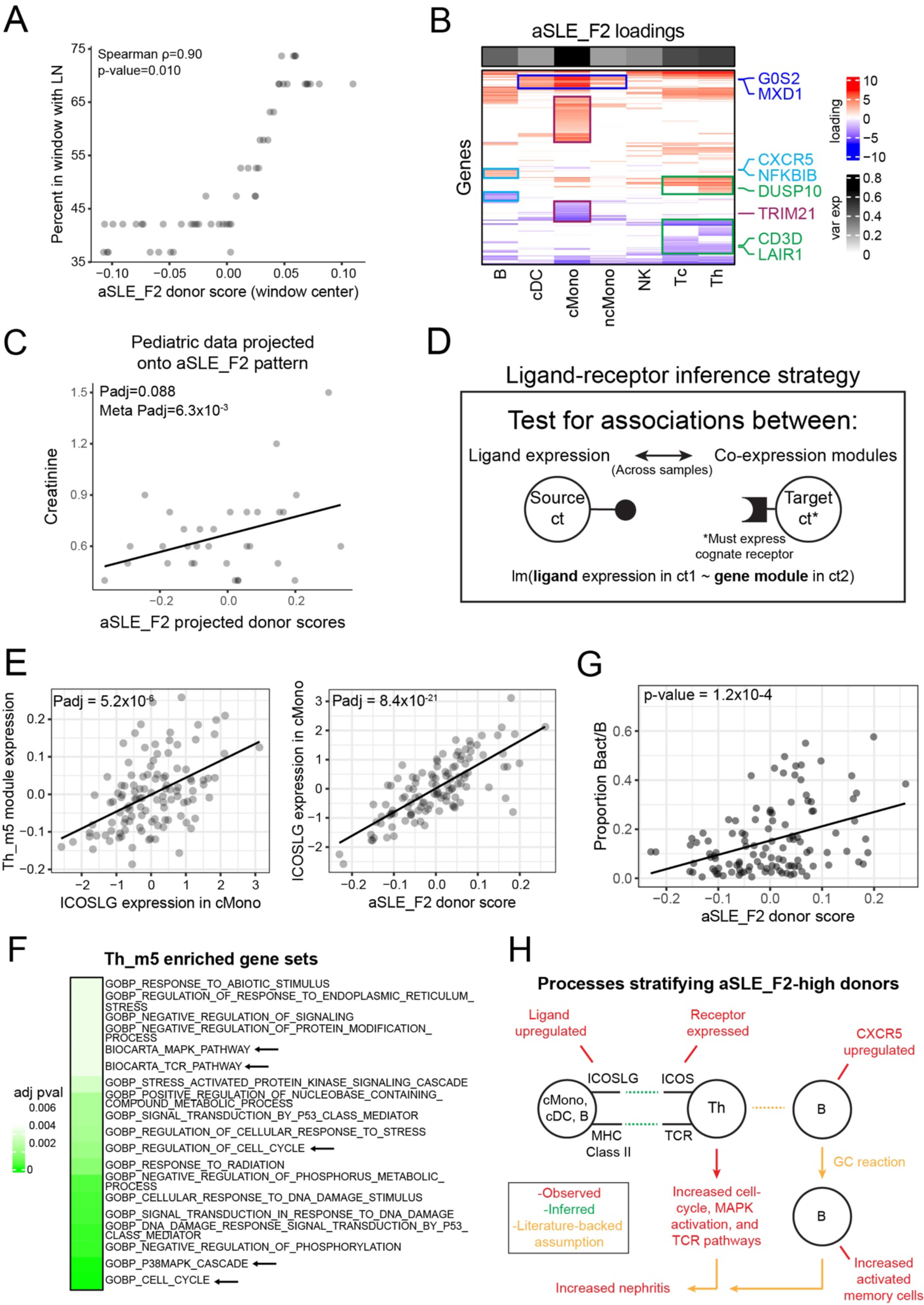
Nephritis-associated factor and ligand-receptor interaction inference. (A) Association between aSLE_F2 donor scores and frequency of lupus nephritis among patients positive for anti-dsDNA autoantibodies. A sliding window was used to compute the percent of patients within the window that had lupus nephritis (Methods). Each point represents the sliding window center. (B) Gene loadings heatmap for aSLE_F3, showing significant genes only. A few relevant genes and the cell types where they are dysregulated are also highlighted. (C) Replicating the nephritis associated signal in the pediatric dataset. The pediatric data were projected onto the aSLE_F2 multicellular pattern, and the resulting donor scores were plotted against creatinine levels. The line is a linear model (also applies to E and G). (D) Overview of the ligand-receptor interaction inference strategy. The bottom text describes the linear model used to test whether ligand expression in a source cell type is significantly associated with a co-expression gene module in a receptor-bearing cell type across donors (Methods). (E) Association between gene module Th_m5 and expression of ligand *ICOSLG* in cMonocytes (left). Association between *ICOSLG* expression in cMonocytes with donor scores for factor aSLE_F2 (right). (F) Enriched gene sets in module Th_m5. Adjusted p-values are shown in green boxes. Enrichment was tested using the hypergeometric test with gene sets from GOBP and BioCarta. Only results with adjusted p-values less than 0.005 are shown. (G) Association between the fraction of activated memory B cells and aSLE_F2 scores. (H) Summary of processes upregulated in the donors with high aSLE_F2 donor scores.

### Inference of ligand-receptor (LR) interactions reveals potential mediators of multicellular patterns

We also sought to identify LR interactions that are candidate mediators of the multicellular patterns identified by scITD. Our approach to inferring LR interactions differs from existing approaches in several ways. Most single-cell LR methods identify interactions based on the upregulation of ligands and their cognate receptors in pairs of cell clusters without regard to the sample of origin. Here, we explicitly test for interactions that are differentially active between samples. In such cases, we expect that donor samples with dysregulated expression of a given ligand (in a source cell type) would also have dysregulated expression of potential downstream target genes (in a target cell type). This is tested by computing the association between ligand expression and gene co-expression modules for pairs of cell types (Figure 3D) (Methods). The gene modules are computed separately for each cell type with WGCNA, using the donor pseudobulk expression matrices. More specifically, when the target cell type expresses the ligand’s cognate receptor to some minimal extent, we use a linear model to test for significant associations between ligand expression and the eigengene of each target cell type co-expression module separately. As a note, we only test for such associations where the source cell type is different than the target cell type. Using signaling network data from NicheNet, encoding ligand-target gene regulatory potential, we validated that the gene modules in our resulting hits were enriched for genes we expected to be perturbed by their associated ligands (Figure S4A, S4B, and S4C) (Methods).

There currently exists only one other single-cell tool for computing differential LR interactions across samples, called Differential NicheNet. To evaluate differences in the results of the two methods, we selected the subset of our results associated with aSLE_F2 and compared them to results generated from running Differential NicheNet on this factor (Methods). The Jaccard overlap in these sets of the results was around 0.1, indicating a small, yet significant overlap (p-value<.001) (Figure S4D and S4E). We also compared validation performance of the two methods using a set of 101 ligand treatment DE datasets. Intuitively, our expectation was that if an inferred LR interaction was differentially active across donors, then we would see the experimentally perturbed target genes also differentially expressed across the same donors. Therefore, for each method, we computed LR interactions and selected LR interactions that included ligands present in one of the treatment datasets. Then, we calculated whether the factor-associated genes in the target cell type were enriched for ligand treatment DE genes (Methods). Both methods achieved similar performance; however, our method demonstrated potentially higher sensitivity as it yielded a larger number of results with supporting evidence from the treatment DE data (Figure S4F and S4G).

One of the top candidate interactions that we identified was between the ligand ICOSLG and its receptor, ICOS, from cMonocytes to Th cells. This is manifested by the association between ICOSLG expression in cMonocytes with a gene module, Th_m5, in Th cells (Figure 3E left). The Th_m5 gene module associated with this ligand was enriched for genes involved in T-cell receptor activation, cell cycle, and p38 MAPK pathways (Figure 3F) consistent with the known co-stimulatory role for ICOSLG-ICOS binding in T cell activation.^40,41^ As expected, we also observed significantly higher ICOSLG NicheNet regulatory potential scores for the genes in Th_m5, as these likely represent potential downstream targets of the ligand (Figure S3C). Additionally, *ICOSLG* was upregulated for individuals with high aSLE_F2 donor scores, connecting this inference to the larger multicellular pattern and nephritis (Figure 3E right). These associations were also replicated in the corresponding nephritis-associated factor in the pediatric dataset, pSLE_F2 (Figure S3D). Corroborating our observations, a previous study showed that T cell ICOS stimulation by myeloid cells contributed to the development of lupus nephritis and was mediated by increased T cell survival.^42^ Related to this finding, we also identified an increase in the proportion of activated memory B cells for individuals with high aSLE_F2 scores (Figure 3G and S3E), which previous studies suggest could be a downstream result of the ICOSLG interaction.^41–43^ This association was identified using an automated procedure we developed to systematically test factors for any changes to subcluster composition by iteratively testing multiple clustering resolutions (Methods). We summarize these findings and existing literature connections in a schematic diagram (Figure 3H). This vignette exemplifies how additional insights can be gained by analyzing the co-occurring processes in multiple cell types, allowing us to generate mechanistic hypotheses driving the patient stratification. As the nephritis association and related molecular observations were not made in either of the original studies which generated the aSLE or pSLE datasets, this further highlights the utility of scITD to yield novel, clinically relevant, and robust findings.

When testing the factors for associations with treatment, we identified that aSLE_F3 was strongly associated with both the use and dosage of the corticosteroid prednisone (Figure 4A and 4D). When projecting the pediatric data onto the aSLE_F3 multicellular pattern, we found that this pattern also significantly stratified the pediatric patients taking oral steroids (Figure 4C). The top associated genes for the factor included *TSC22D3*, a known glucocorticoid response gene, which was upregulated in multiple cell types (Figure 4B). This is an expected result, as prednisone binds the glucocorticoid receptor. However, some glucocorticoid response genes such as *DUSP1*, were found to only be upregulated in T cells. Similarly, GSEA analysis for this factor confirmed the expected enrichment of hormone response genes in all cell types except B cells and, specifically, corticosteroid response genes in Tc, Th, and cDC cells (Padj<.05).

**Figure 4.**
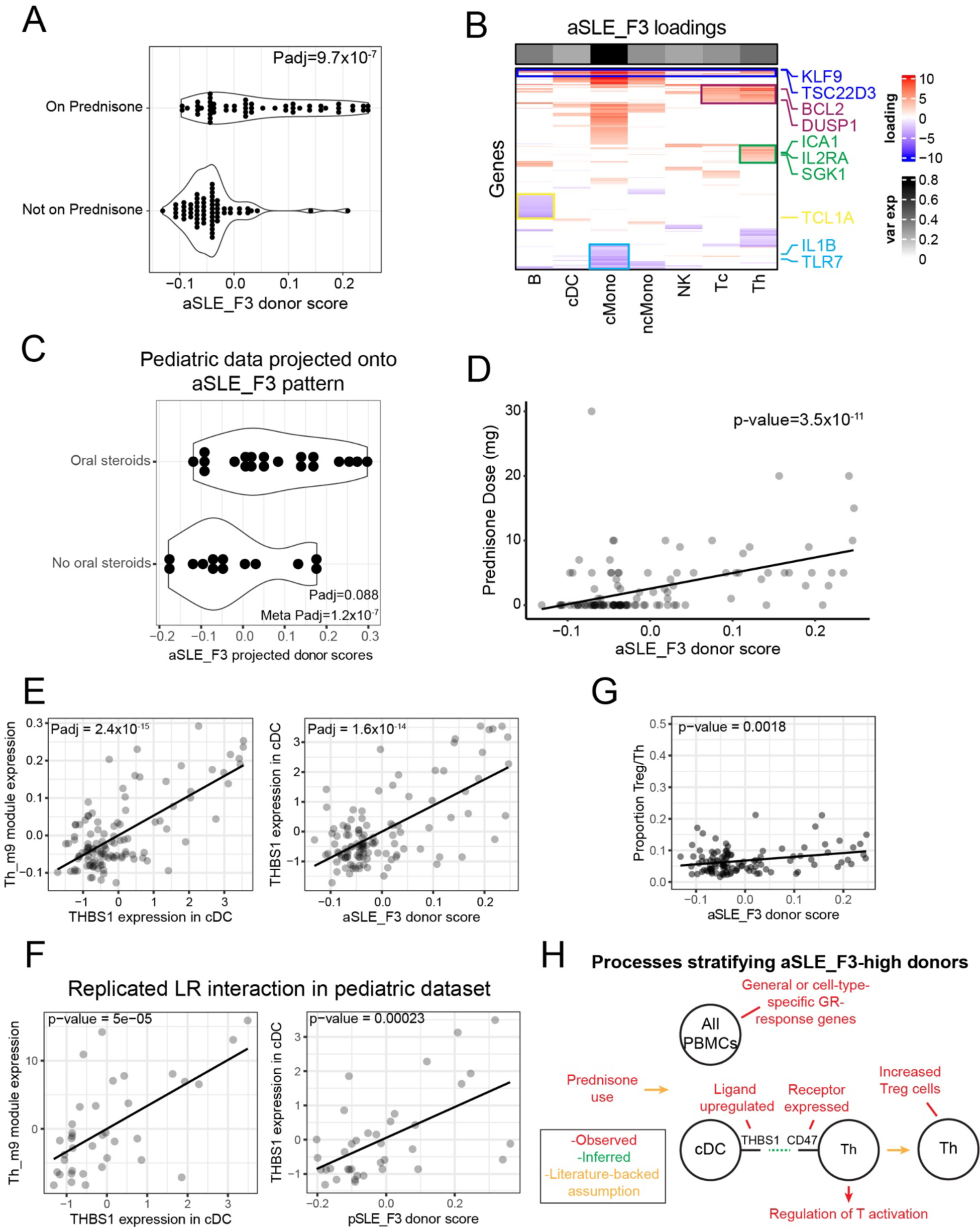
scITD identifies a prednisone-associated factor with cell-type-specific effects. (A) Association between aSLE_F3 and prednisone use. The significance of each association was computed using logistic regression with a likelihood-ratio test. (B) Gene loadings heatmap for aSLE_F3, showing significant genes only. A few relevant genes and the cell types where they are dysregulated are also highlighted. (C) Replicating the prednisone associated signal in the pediatric dataset. The pediatric data were projected onto the aSLE_F3 multicellular pattern and the resulting donor scores were plotted against oral steroid use metadata. Significance was evaluated using logistic regression with a likelihood-ratio test. (D) Association between aSLE_F3 and prednisone dose. The outlier with the highest prednisone dose was not included in the calculation of the linear model p-value but is still shown in the plot. The p-value was calculated using a linear model F-test. The line is a linear model and significance was evaluated with an F-test (also applies to E, F, and G). (E) Association between gene module Th_m9 and expression of ligand *THBS1* in cDCs (left). Association between *THBS1* expression in cDCs with donor scores for factor aSLE_F3 (right). (F) Replicating the THBS1 LR interaction in the pediatric dataset with the corresponding pediatric factor, pSLE_F3. The same genes from the Th-m9 module of the adult SLE dataset were used to compute the gene module scores. (G) Association between the fraction of Treg cells and aSLE_F3 scores. (H) Summary of processes upregulated in the donors with high aSLE_F3 donor scores.

Our LR inference approach also identified several strong candidate interactions where the ligand’s expression was associated with aSLE_F3. One such candidate included the ligand *THBS1* in cDCs which positively correlated with the Th_m9 co-expression module and aSLE_F3 (Figure 4E). As with aSLE_F3, module Th_m9 was significantly enriched (adjusted p-values < 0.01) for genes involved in response to hormone (e.g., *KLF9*, *TXNIP*) and regulation of T cell activation (e.g., SOCS1, *NFKBIZ*). We also observed significantly higher *THBS1* NicheNet regulatory potential scores for genes in the Th_m9 gene module compared to genes in all other modules (p-value = 2.8×10^-8^), supporting the inference of this interaction. We were further able to reproduce the evidence for this interaction in the corresponding factor of the pediatric decomposition, pSLE_F3 (Figure 4F). Previous studies have also shown that dendritic cell-derived *THBS1* can promote Treg development when interacting with integrin-associated protein (CD47).^44^ Therefore, we tested for an association between the aSLE_F3 donor scores and Treg proportions. This association was statistically significant, with a relative expansion of this Th subpopulation in the prednisone-taking patients (Figure 4G). Unlike the aSLE_F1-high donors, however, the aSLE_F3-high patients did not have increased ISG expression. These results highlight a high confidence shift in intercellular communication upon prednisone use that may mechanistically contribute to its anti-inflammatory effects. A summary of the biological findings linked to this factor are summarized in a schematic diagram (Figure 4H).

### Differences between corresponding adult and pediatric SLE factors

As mentioned above, applying scITD to the pediatric SLE dataset yielded 3 factors that stratified patients in a highly similar manner to adult SLE factors. Therefore, we sought to test whether there were any meaningful differences between the corresponding factors. We started by comparing the gene-factor associations between the datasets (Figure S5A). While generally highly correlated, we posited that there may still be subtle differences in the genes involved in each dataset’s factors. Therefore, for each set of corresponding factors, we computed the difference in each gene’s factor association coefficient between datasets. Dataset-specific genes are expected to exhibit higher association differences (Figure S5B). We then used these difference values in GSEA to identify dataset differences at the gene set level (Methods). We checked that this procedure did not produce excess false positives in a null comparison that consisted of the adult SLE dataset and the same dataset downsampled to 90% of donors (Figure S5F). In comparing aSLE_F1 with pSLE_F1 (the corresponding ISG factors), we identified significantly enriched gene sets in every cell type, pointing to systematic differences in the interferon-response between these two populations of SLE patients (Figure S5C). Some of these differences included activation, differentiation, and migration gene sets in the lymphocytes as well as cell-cycle gene sets in the myeloid cells (Figure S5C). The directionality of the leading-edge genes for these gene sets suggests greater immunologic activity associated with ISGs in the pediatric patients compared to the adults (Figure S5D). For example, *BATF* (mediates class-switch recombination) in B cells and PRF1 (involved in cell killing) in Tc cells had larger linear model associations with pSLE_F1 compared to aSLE_F1. This could indicate that interferon has a differential impact on lymphocytes in pediatric compared to adult cases, with certain immunologic functions enhanced in pediatric cases. In our comparison of the nephritis factor, aSLE_F2, and its corresponding pediatric factor, pSLE_F2, we identified differences in Tc cells with regard to activation and differentiation related genes (Figure S5E top). These differences could potentially point to reasons why pediatric SLE cases are more likely to develop renal manifestations or why pediatric cases have different incidence of nephritis subtypes compared to adult cases.^45^ We also compared the corresponding prednisone-associated factors, aSLE_F3 and pSLE_F3, though only finding weak enrichments for various immune-related gene sets (Figure S5E bottom). In summary, while the larger multicellular patterns are similar across datasets, there exist more nuanced, biologically relevant differences that distinguish them.

### Multicellular expression patterns stratifying COVID-19 patients include disease-specific patterns and conserved immune regulation also found in SLE

Given the prominent type-1 ISG signal in SLE patients, a gene signature normally observed in accute viral infection, we next applied scITD to analyze a large scRNA-seq dataset consisting of 83 COVID-19 patients and 20 healthy controls.^46^ The patients demonstrated varying degrees of disease severity at the time of sample collection, ranging from asymptomatic to critical. We again limited our analysis to the major cell populations (Figure 5A). We ran scITD on this dataset to extract 9 factors, labeled CVD_F1, CVD_F2, etc. (Figure 5B). Surprisingly, 5 of the aSLE factors stratified the COVID-19 patients in a highly similar manner to 5 of these factors (Figure 5B and 5C). For example, we observed that CVD_F1 (which explains most variance in the dataset) stratified samples in much the same way as aSLE_F1, seen by the high correlation in donor scores (Figure 5B). As with aSLE_F1, CVD was characterized by pan-cell type ISG expression (Figure 5D). Since aSLE_F1 was associated with SLE disease activity, we tested whether CVD_F1 was associated with disease severity. We did not observe a significant association between CVD_F1 donor scores and the fine-grained severity category using a simple linear model (Figure 5B bottom). However, we noticed that both healthy donors and COVID-19 critical patients had significantly lower CVD_F1 donor scores (by an enrichment test) compared to the non-critical COVID-19 patients (Figure 5E). These results are consistent with the known effect of autoantibodies against type-1 interferons as a major determinant of critical COVID-19.^47,16^ To address the possibility that this observation was due to the CVD_F1 correlation with age, we regressed age out of the donor score and used the residuals in the same enrichment test, still yielding significant enrichments for the COVID-19 categories (Padj_critical_=0.0038, Padj_non-critical_=0.00057, and Padj_healthy_=0.16).

**Figure 5.**
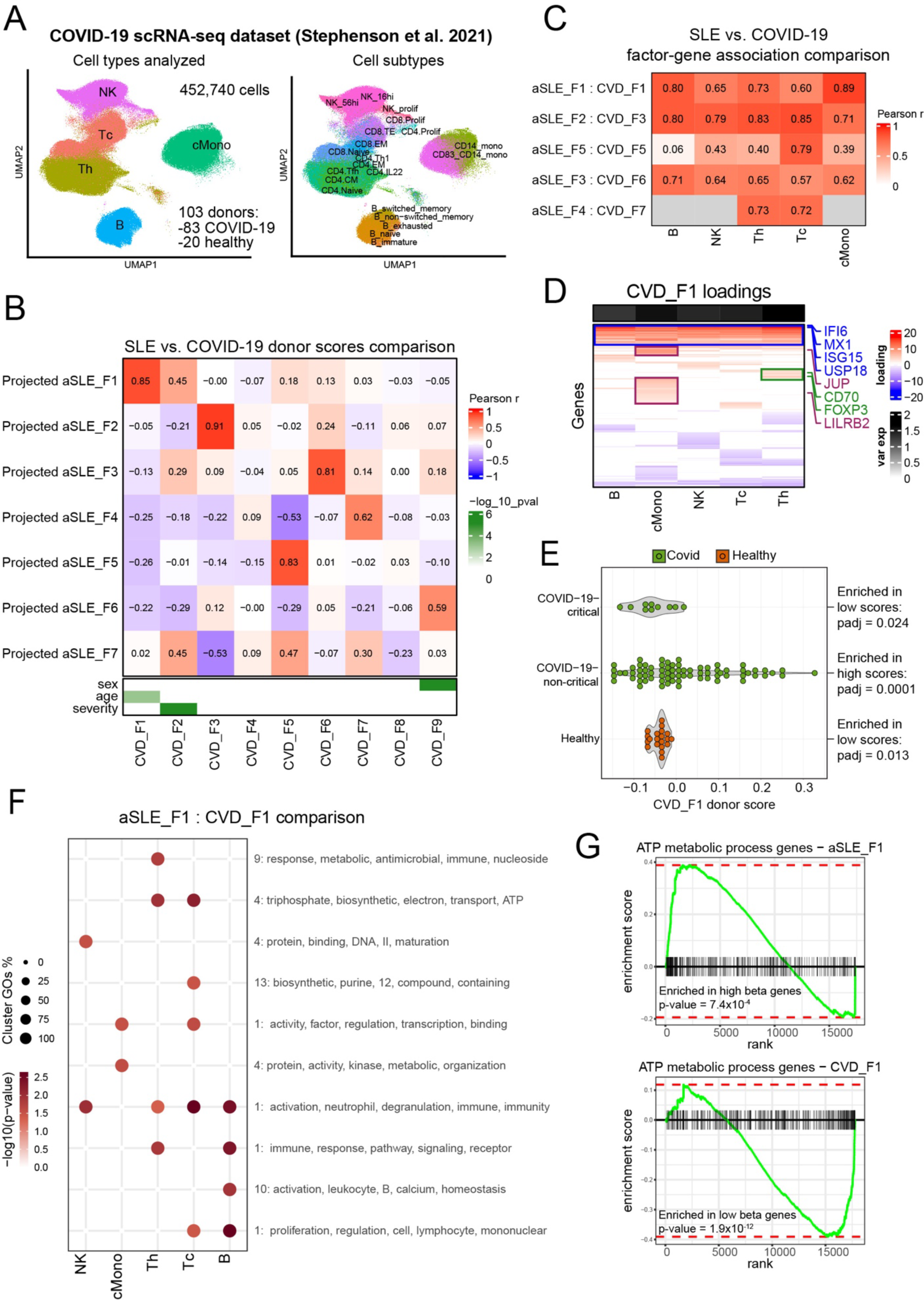
Multicellular patterns that stratify SLE patients also stratify patients in a COVID-19 dataset. (A) UMAP plot of single-cell gene expression from Stephenson et al. colored by the major cell types used in the scITD analysis (left) or cell subtype annotations (right). (B) Correlations between projected donor scores (using aSLE factors) and COVID-19 donor scores. Metadata associations with COVID-19 factors are shown in the bottom heatmap. Association p-values were calculated using univariate linear model F-tests. (C) Pearson correlations of gene-factor association coefficients between aSLE and CVD datasets. Only genes that had significant associations with a given factor in either dataset were used for computing the correlations. Gray color indicates cell types that were not tested because they had fewer than 10 significant genes total. (D) Gene loadings heatmap for aSLE_F3, showing significant genes only. A few relevant genes and the cell types where they are dysregulated are also highlighted. (E) Association between COVID-19 status and CVD_F1 donor scores. FGSEA running enrichment tests were used to calculate enrichment of patient groups at either end of the CVD_F1 donor scores. (F) Gene set enrichment analysis using dataset-specific upregulated or downregulated genes when comparing aSLE_F1 and CVD_F1. Gene sets were collapsed in two stages, first by overlapping sets of genes and then by the pattern of gene sets across cell types. The number in front of the gene set terms indicates the number of sets of gene sets with similar terms that clustered together (Methods). (A) Differential enrichment of ATP metabolic process genes in Tc cells along either aSLE_F1 (top) or CVD_F1 (bottom). FGSEA was used to calculate p-values.

As ISG appears to play opposite roles in COVID-19 (protective role) compared to SLE (pathogenic role), we aimed to identify differences in the ISG expression patterns between acute viral infection and autoimmunity. By applying the same strategy used to compare corresponding adult and pediatric factors, we identified multiple enriched gene sets among the genes that were differential between aSLE_F1 and CVD_F1 (Figure 5F). These included ATP metabolic process genes (T cells) and activation/proliferation genes (multiple cell types) among others. The SLE-specific enrichment of ATP metabolic process genes in ISG-high Tc cells (Figure 5G) is particularly intriguing as one previous study demonstrated striking metabolic changes in Tc cells that take place only in response to chronic interferon exposure and were absent with short-term exposure.^48^ This is in line with our expectation that interferon stimulation in SLE patients is of a chronic nature whereas in COVID-19 it is acute.

Given that the aSLE and COVID-19 datasets had different distributions of donor age and sex (Figure S6A), we tested whether these could be responsible for the observed differences in the corresponding ISG patterns, aSLE_F1 and CVD_F1. We hypothesized that if age and sex differences were driving the differences between these two factors, then we would see the gene-factor association coefficient deltas shrink to 0 after balancing the datasets for age and sex. Instead, we observed no obvious biasing in these statistics after balancing the datasets (Figure S6B, S6C, and S6D). Importantly, the leading-edge genes corresponding to the enriched gene sets in Figure 5F were also unaffected (Figure S6D).

Next, we further investigated factor CVD_F2, which was unique to the COVID-19 dataset in that it did not highly correlate with any SLE factors (Figure 5B). We further observed that CVD_F2 was associated with COVID-19 severity in a more continuous manner, stratifying donors from healthy to critical disease status linearly along the factor (Figure 6A). The multicellular pattern for this factor consisted of multiple cell-type-specific biological processes including cell-cycle (NK and T cells) and various signaling cascades (in cMonocytes, NK, and T cells) among others (Figure 6B). We aimed to replicate this severity-associated multicellular pattern by analyzing another COVID-19 scRNA-seq dataset. The validation dataset from van der Wijst et al.^16^ consisted of a smaller number of donors but included the same cell types and a similar range of disease severity (not including asymptomatic or mild cases). We projected the validation dataset onto the CVD_F2 multicellular pattern and similarly found that this pattern significantly stratified patients by disease severity (Figure 6C). A meta-analysis combining the results from these two datasets yielded a Fisher p-value of 4.3×10^-17^ (P_CVD_ = 5.2×10^-11^, P_VAL_ = 2.0×10^-8^).

**Figure 6.**
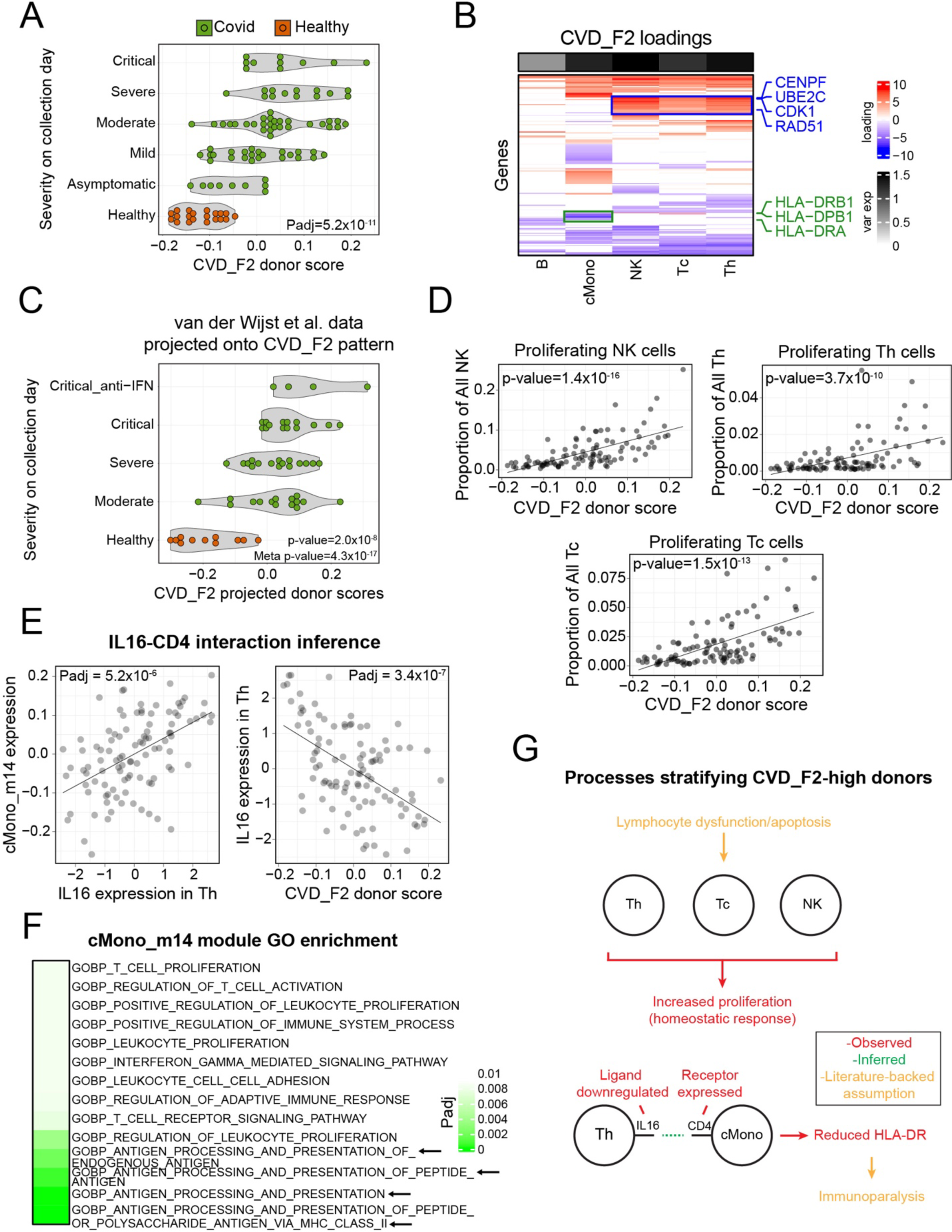
Analysis of a COVID-19 severity associated multicellular pattern. (A) Association between COVID-19 severity at sample collection and CVD_F2 donor scores from the decomposition of the Stephenson et al. COVID-19 dataset. The significance of the association was calculated with a linear model F-test. (B) Gene loadings heatmap for CVD_F2, showing significant genes only. A few relevant genes and the cell types where they are dysregulated are also highlighted. (C) Replicating the COVID-19 severity association in the van der Wijst et al. validation dataset. The validation data were projected onto the CVD_F2 multicellular pattern, and the resulting donor scores were plotted against COVID-19 severity metadata. The association p-value was calculated the same way as in (A). The color legend is also the same as in (A). (D) CVD_F2 associations with cell subtype proportions for proliferating T cell and NK cell populations. The p-values were calculated using linear model F-tests. (E) A potential LR interaction between IL16 and CD4 (Th cells to monocytes) identified in the Stephenson et al. dataset. The left plot shows the association of gene module cMono_m14 with expression of ligand *IL16* in Th cells. The plot on the right shows the association of *IL16* expression in Th cells with donor scores for CVD_F2. The line is a linear model. (F) Top enriched GO gene sets in co-expression module cMono_m14 from the Stephenson et al. dataset (adjusted p-values < 0.009). (G) Summary of processes upregulated in the donors with high CVD_F2 donor scores.

We further examined this factor for associations with cell subcluster proportions using subcluster annotations from the original COVID-19 study that generated this dataset. We found that the more severe patients had reduced proportions of activated Th cells (IL22+) as well as increased proportions of terminal effector Tc cells (Figure S7A). These 2 subcluster proportion associations were also reported in the original study that generated this dataset.^46^ However, by using scITD we found that the CVD_F2-high donors had significantly increased proportions of proliferating Th cells, Tc cells, and NK cells (Figure 6D), consistent with the proliferation markers highlighted in the loadings matrix (Figure 6B). By further applying our LR inference technique, we identified a strong candidate LR interaction connected to this multicellular pattern. The interaction included the ligand IL16 expressed from Th cells interacting with the CD4 receptor on cMonocytes. Specifically, we observed high correlations between *IL16* expression in Th cells and various co-expression gene modules in cMonocytes, including cMono_m14 (Figure 6E left). Module cMono_m14 was significantly enriched for MHC Class II genes (Figure 6F), indicating a possible role of Th derived *IL16* in regulating the expression of these genes in monocytes. Corroborating this, earlier studies have shown that IL16 can upregulate monocytic MHC Class II genes, including *HLA-DR*.^49^ Here, we further found Th *IL16* to be downregulated in donors with high CVD_F2 donor scores (Figure 6E right), and the CVD_F2 multicellular pattern shows that these donors also have downregulation of *HLA-DR* expression in monocytes (Figure 6B). Interestingly, reduced expression of *HLA-DR* in monocytes has also been found in patients with sepsis, a process that shares many of the same pathophysiological features as severe COVID-19.^50,51^ Two recent studies have also identified reduced levels of HLA-DR protein on monocytes of critically ill COVID-19 patients.^52,53^ Given our result and the prior literature, further studies should be conducted to determine whether reduced *IL16* contributes to increased COVID-19 severity via reduced *HLA-DR* expression. Lastly, the severe and critical patients in the validation dataset also had significantly reduced *IL16* expression in their Th cells (Figure S7C right) matching our observation from the Stephenson et al. data. Similarly, *IL16* was again positively associated with a co-expression module in cMonocytes (Figure S7C left) that was enriched for MHC Class II genes (Figure S7D). The consistency of these results across independently generated datasets bolsters our confidence in the connection between this multicellular pattern and COVID-19 disease severity and highlights the conserved nature of a potentially relevant cell-cell interaction.

## Discussion

We developed a novel computational tool for stratifying healthy or disease individuals using scRNA-seq data. Uniquely, our method, called scITD, extracts recurrent patterns of gene expression involving one or more cell types that vary across individuals. This is useful because tissue biology and its dysregulation often involves coordinated changes to multiple cell types. We principally applied our tool to analyze several large scRNA-seq dataset of PBMCs from individuals with SLE, identifying multicellular patterns of transcriptional variation that stratified patients. This analysis yielded both expected patterns (e.g., ISG expression across multiple cell types) and new ones linked to various phenotypes (e.g., nephritis, prednisone use, age, ethnicity, etc.). By further analyzing the cell-type-specific biological processes and the cell-cell interactions that made up each multicellular pattern, we illustrated how scITD can be used to gain a more detailed understanding of the cellular components relevant to a given disease, their connectivity and links to the associated phenotypes.

Analysis of the second, pediatric SLE dataset reproduced the key patterns and associations observed in adult patients, highlighting the robustness of these signals. This indicated that main sources of transcriptional variation across patients were conserved in both patient populations. However, in comparing cognate factors between the two datasets, we found subtle differences that may partially explain the phenotypic differences between adult and pediatric SLE. We further applied scITD to stratify donors in a COVID-19 scRNA-seq dataset. Surprisingly, this also revealed several multicellular patterns that were highly correlated with those that stratified the SLE patients. In comparing the cognate ISG-factors between datasets, we observed differences that can potentially be attributed to the chronic (SLE) versus acute (COVID-19) nature of interferon stimulation in each disease. Despite these subtle differences, we believe this high degree of conservation points to there being a limited number of stable states that a cell system can take on. Genetics can likely play a role in reshaping these states and could be responsible for the differences we observed when comparing corresponding factors across datasets. We also found a multicellular pattern unique to the COVID-19 data, which was associated with a continuum of disease severity. Such patterns unique to a given disease may highlight molecular states linked to fundamentally different types of phenotypes, such as a septic shock-like response seen in a subset of critical covid patients but not in any SLE patients. These results highlight the need to generate datasets sampling individuals with many different conditions, so that we can characterize all common configurations of covarying immune cell states.

In addition to studying inter-individual variation, scITD may also be informative in other types of analyses. For one, scITD can be applied to investigate how different technologies, processing techniques, and disease models impact expression jointly within each cell type. We briefly demonstrated how scITD can be used in this way to better understand batch effects (Supplementary Note S2). More work in this area could lead to better-informed designs of batch-correction methods tailored for various scenarios. Another use case for the scITD is in the study of multi-tissue patterns of gene expression (as opposed to multi-cell type patterns). Since scITD uses scRNA-seq data at the pseudobulk-level, the tool can be directly applied to bulk RNA-seq datasets generated from multiple donors with multiple tissue types (e.g., the GTEx studies). This could allow one to connect gene expression changes in the blood with the expression states of less accessible tissues. Overall, scITD enables researchers to analyze gene expression covariation among different cell types, extending our ability to study the complex biological processes that stratify individuals in health and disease.

### Limitations of the Study

A notable limitation of our approach is that it does not reveal causal relationships that may be underlying multi-cellular patterns of transcriptional variation. In interpreting the results of the current study, we relied on corroborative evidence from scientific literature or information from signaling networks. We hope that future iterations of our approach will enable the identification of potentially causal events by analyzing cis-regulating genetic variants using Mendelian randomization or other techniques. Furthermore, since the presented method is unsupervised, it can only reveal patterns that exhibit sufficiently high variation across the examined samples. In that regard, the approach may fail to separate factors that would stratify individuals along a desired variable of interest (e.g., disease status), unless there is substantial expression variation along that axis. We hope to add options for carrying out semi-supervised decomposition in the future, which would allow to extract one or more factors stratifying patients along specific variables of interest. Given linear assumptions of the employed tensor decomposition procedure, the approach will have difficulty capturing highly non-linear processes in isolated factors. Finally, deciding on the appropriate number of factors to extract remains challenging. We have implemented a strategy to do so by monitoring diminishing amount of variance explained by each successive factor, but such progressions may not yield a clear threshold.

## Methods

### scITD pipeline and details

#### Data preprocessing

Before running the tensor decomposition, it is crucial to first generate the tensor with dimensions donors by genes by cell types (*N* × *G* × *C*). This data structure is generated by forming a donor-by-gene pseudobulk expression matrix (dimensions *N* × *G*) for each cell type separately. These matrices are then stacked together along the 3^rd^ dimension to form the tensor. To generate each donor-by-gene pseudobulk expression matrix, we first subset the gene-by-cells UMI counts matrix by a cell type. Then, for each gene, we sum counts across cells belonging to the same donors. Then, for each donor, we sum all counts across gene to get the donor’s library size for the given cell type. Then, we apply the trimmed-mean of M values (TMM) method in edgeR^54^ to adjust library sizes of the pseudobulked counts. We normalize the pseudobulk counts by dividing by each donor’s adjusted library size, multiplying by a scale factor (typically 10000), and applying a log-transformation with a pseudo-count of 1. Before this previous step, we only retain donors that have at least a minimum number of cells in every cell type, as the Tucker decomposition does not allow for NA values in the tensor. As with PCA, it is helpful to reduce the genes included to those that are highly variable across donors. Therefore, we compute the normalized variance for each gene in each cell type, using the method from pagoda2.^55^ However, in order to stack the donor-by-gene matrices together to form the tensor, they must each contain the same genes. Therefore, we select the top overdispersed genes from each donor-by-gene matrix and subset these matrices to the union of overdispersed genes from all cell types. Next, for each donor-by-gene matrix, normalized gene expression values are centered and unit scaled across donors. This forces each gene to have the same average pseudobulk expression (i.e., 0) in each cell type, ensuring that cell type differences in baseline expression will no longer be an axis of variation in the data (i.e., only inter-donor variation remains). We then rescale variance for each gene by its normalized variance. This allows genes with greater or less biological variation to contribute proportionally to the decomposition. This is done by multiplying the expression by the normalized variance value to some power. The power should be set to 0.5 for the resulting variance to equal the normalized variance. We note that increasing the value of the power slightly (often between 1-2) can sometimes improve the quality of the decomposition. Finally, the pseudobulked donor-by-gene matrices from all cell types are stacked together to form the tensor.

#### Tensor decomposition

Next, we apply the Tucker tensor decomposition to the preprocessed pseudobulk tensor. For this, we use the R package, rTensor,^56^ which implements Higher-Order Orthogonal Iteration (HOOI) to compute the Tucker decomposition. We chose this tensor decomposition over others because of its rotational flexibility, which can help with interpretability, as well as its relatively small number of estimated parameters. The formal algorithmic procedure for HOOI can be viewed in the following publication.^57^ HOOI outputs a separate factor matrix for each mode of the tensor (i.e. donor, gene, or cell type dimension) and a core tensor that can be multiplied together to reconstruct the approximation of the starting tensor. The matrices for each mode are allowed to have differing numbers of factors, and therefore from here on we specify these as “donor factors”, “gene factors”, and “cell type factors”, respectively. Thus, the three separate factor matrices are of dimensions donors-by-donor factors, genes-by-gene factors, and cell-types-by-cell-type factors. The core tensor is of dimensions donor factors-by-gene factors-by-cell-type factors. The standard data reconstruction using these objects is as follows:

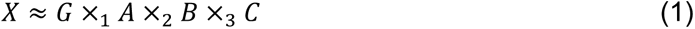

Here, *X* is the reconstructed tensor, *A* is the donor factor matrix, *B* is the gene factor matrix, *C* is the cell type factor matrix, and *G* is the core tensor. The operator ×_$_ indicates multiplication of the matrix on the right side of the operator by the tensor on the left side of the operator along the n^th^ mode of the tensor, also called the n-mode product.^58^

As the factor matrices and core tensor are challenging to interpret on their own, we rearrange and combine the terms to form a structure that looks more similar to that of a standard matrix decomposition. Such a structure would have only one set of factors, where each factor has a component for samples (i.e. scores) and a component for the variables (i.e. loadings). From our decomposition output, we can directly consider our donor-by-donor factor matrix to be our matrix of sample scores, but it is necessary to rearrange and combine the other terms to form the loadings. To obtain this, we multiply the core tensor by the gene factor matrix and cell type factor matrix using n-mode product. This yields a new tensor of dimensions donor factors-by-genes-by-cell types. This is the object we consider to be the loadings tensor, where each factor slice is a multicellular expression pattern (dimensions *G* × *C*) (Figure S1A). This reordering of the terms in the product is valid because the order of multiplication does not matter when reconstructing the data, as long as the multiplication mode also changes accordingly.^58^ This reordering and simplification are done as follows:

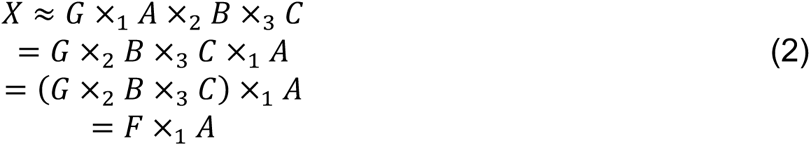

Here, *F* is the loadings tensor of dimensions donor factors-by-genes-by-cell types, the rest of the terms are the same as in the previous equation. As a note, in Figure S1A this reconstruction is shown backwards as *A* ×_1_ *F* simply to demonstrate how the tensor times matrix multiplication yields the reconstructed tensor of correct dimensions, even though convention is to write the multiplication in reverse.

#### Factor rotations

One unique attribute of the Tucker tensor decomposition is that the factors can be rotated to improve their interpretability.^59,60^ We explored the practical implications of two different rotations that users should consider when applying scITD to their own data (Supplementary Note S1). When designing the rotations, we leverage the fact that the factor matrices from the output of the Tucker decomposition can be rotated by any non-singular square matrix as long as the core tensor is counter-rotated by the inverse of the rotation matrix. The counter-rotation ensures that the reconstruction error will remain unchanged. The core tensor can also be rotated similarly as long as the factor matrices are counter-rotated accordingly. Here, we explored the use of various rotations. One such approach we tested was applying ICA rotation to the donor scores matrix, and counter-rotating the core tensor before computing the loadings tensor. As a note, we needed to normalize the rotated donor scores matrix because ICA does not preserve lengths. After rotating the donor scores matrix, the core tensor is counter-rotated as follows:

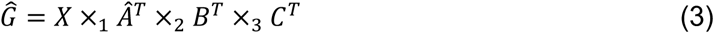

Here, *Ĝ* is the new core tensor, *Â*^*T*^ is the transpose of the ICA rotated (normalized) donor factor matrix, *C*^*T*^ is the transpose of the cell type factor matrix, *B*^*T*^ is the transpose of the gene factor matrix. Then, we substitute *Ĝ*^T^ for *G* when calculating the loadings tensor *F* in equation 2.

However, this first approach often made it challenging to interpret the multicellular expression patterns, as the loadings were often correlated across factors (Supplementary Note S1). Therefore, we tested a second type of rotation, with the goal of optimizing independence of the loadings instead of donor scores. This approach involves a two-step rotation procedure which is a hybrid of ICA and varimax applied to the distinct terms that make up the loadings tensor. The intuition behind this approach is to create a core tensor, where each donor factor represents some combination of biologically distinct gene sets (gene factors) in the different cell types. This is achieved by rotating donor factors of the core tensor to a simple structure. This helps to ensure that each gene factor only partakes primarily in one donor factor. This can be achieved by applying varimax to the core tensor. However, it is also necessary to ensure that all gene factors are independent of one another. Otherwise, the optimized core tensor may still yield donor factor loadings with similar sets of relevant genes. To accomplish this, we apply rotations in two separate steps. In the first step, we apply ICA to the gene factor matrix and counter-rotate the core tensor (equation 4A). In the second step, we optimize the core tensor by the varimax rotation and counter-rotate the donor matrix (equation 4B). This is calculated as follows:

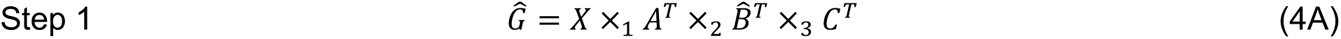

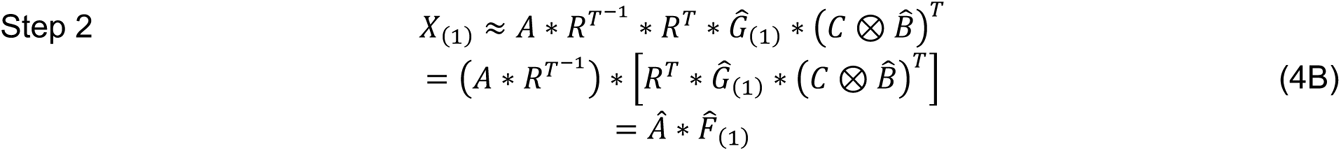

Here, all variables are as previously described, with the addition of *B̂*, which represents the ICA rotated gene factor matrix, and *R* which now represents the orthonormal rotation matrix found by varimax when applied to 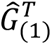 (the subscript indicates the mode along which the tensor is unfolded). The symbol ⊗ is the Kronecker product. For this approach, we use the identity matrix for *C*. For all of our analyses, we also set the ranks of *C* equal to the number of cell types, since we used a relatively small number of cell types. This is the primary rotation used throughout the paper, with the exception of specific cases where we expected to extract discrete sample groupings such as in the in vitro treatment data (Figure 1B) and batch-effect analysis (Supplementary Note 2).

#### Projecting new data onto factors

Before projecting new data onto existing factors, we recompute the original loadings without the core rotation as *F*_(1)_ = *Ĝ*_(1)_ * (*C* ⊗ *B̂*)^*T*^. Then, we calculate the new donor scores by 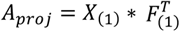. Lastly, the columns of the new scores are normalized to a magnitude of 1 and are rotated by applying the varimax rotation matrix from the core optimization as *Â*_*proj*_ = *A_*proj*_ * *R*^*T*-1^*.

#### Explained variance calculation

For visualization purposes, we order the factors in the donor scores matrix from highest to lowest explained variance. To calculate the variance explained by a given factor, we first compute the reconstructed tensor *X̃* using only the selected factor. Then variance explained is simply the calculation for the coefficient of variation:

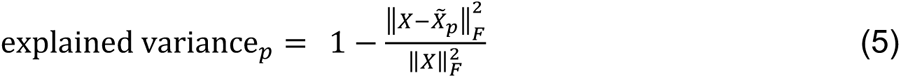

The subscript *p* indicates the factor used for reconstructing the data. The subscript *F* on the double brackets indicates the Frobenius norm. In the loadings matrices, we also display the amount of variance explained by each cell type component. To compute explained variance for individual cell type components within a factor, all values of the reconstructed tensor for all other cell types not under consideration are set to 0:

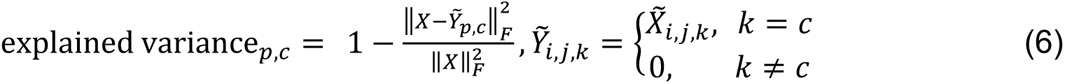

The subscript *c* refers to the cell type being used for calculating explained variance. The subscripts *i*, *j*, *k* refer to the donor, gene, and cell type index of the tensor, respectively.

### Simulation study

To generate the simulations we used the R package, Splatter.^61^ Specifically, we applied the Splatter to generate four subpopulations for each of the two cell types (Figure 2A). To generate a set of four subpopulations, two separate group simulations were run and concatenated together. Each simulation generated two groups of cells separated by some DE genes. Half of the cells from the first population of the first simulation were matched to those of the first population in the second simulation. Likewise, half of the cells from the second population of the first simulation were matched to the other half of the cells from the first population of the second simulation and so on. We then assigned groups of donors (by those with upregulation of each multicellular pattern) randomly to cells from the specified cell subpopulations.

We ran the data through the standard scITD pipeline. However, in this analysis, we did not reduce the tensor to only the most variable genes, so that it would be possible to use all genes in the AUC calculation. We computed the Tucker decomposition to two donor factors and four gene factors, as suggested by our rank determination method. We then calculated gene significance p-values for each cell type in each factor. This was done using linear model F-tests with expression as the explanatory variable and donor scores as the response variable. This technique is also used in the main dataset analyses below to identify significant genes per factor. Then, we used these p-values to calculate the ROC AUC for predicting ground truth DE genes that distinguish each of the two multicellular patterns. Lastly, we subsampled the simulated dataset to varying sizes to determine the robustness of our method to a reduced signal-to-noise ratio.

### Perez et al. SLE dataset processing

The SLE scRNA-seq dataset was originally demultiplexed using an updated version of *demuxlet*,^3^ and quality control measures were applied using Scanpy^62^ with the default parameters. We further filtered out cells with over 10% of their UMIs attributed to mitochondrial genes. This dataset originally contained over 200 donors with transcriptomes from over 1.2 million cells. To make it possible to use this dataset with scITD in its current framework, we reduced the dataset down to one sample per donor, used only the largest cell clusters, and restricted it to only those donors with at least 20 cells in each major cell cluster. Cell clusters and annotations from the original study were used. This left us with 171 donors and 632,733 cells. The median number of cells per donor for B, NK, Th, Tc, cDC, cMonocytes, and ncMonocytes were 421, 264, 1145, 688, 50, 939, and 144 cells respectively. According to our simulation study discussed previously, these quantities were more than sufficient to extract multicellular patterns with high accuracy. We formed the expression tensor using the standard scITD pipeline, and we also applied ComBat batch correction^63^ at the level of 10X lanes to each cell-type slice of the tensor. For our primary SLE analysis, we used the hybrid rotation method and decomposed the data into 7 donor factors and 20 gene factors. We used the same ranks parameters for the decomposition run on only the SLE donors. We computed associations between the donor scores for each factor and the metadata variables and 41 clinical features. The statistical tests used depended on the type of the variable being tested. For the ordinal variables such as SLEDAI score and SLICC score, we employed the ordinal logistic regression with the “probit” method, and p-values were calculated using the resulting t-statistics. For binary variables such as the presence of symptoms or prednisone use, we employed a logistic regression with a chi-square test for significance. For continuous variables we used linear model F-tests. We only tested the binary variables that were present in at least 20 donors. Multiple hypothesis test correction was applied with the BH procedure. To test for the association of aSLE_F2 with the co-occurrence of lupus nephritis and anti-dsDNA autoantibodies, we first removed any donor scores for donors that did not have anti-dsDNA autoantibodies present. Then we used a sliding window of size 19 to calculate the number of donors within the window that also had lupus nephritis. To calculate a p-value we randomly shuffled donor scores and computed the Spearman correlation between the donor scores and the sliding window count. We repeated this procedure 10000 times to generate a null distribution of correlation values. The null distribution was then used to calculate a p-value, by counting the number of null instances with a larger correlation than the one we observed with the unshuffled data.

### IFN-beta experiment data processing

We only kept the cell barcodes labeled as singlets and used the cell-type annotations ascribed by the authors who generated the data.^3^ DE analysis results were also used directly from the original paper where the data were generated. We ran the Tucker decomposition to extract two donor factors, four gene factors. We used the ICA rotation method on donor scores for this dataset, although the results were practically identical when obtained using the other rotations as well. The downsampling procedure was performed as described above with the simulated data, except that instead of AUC, we report the Spearman correlation between the loadings and log2FC values from the DE analysis.

### Procedure for rank determination

To help determine the appropriate ranks to decompose the tensor to, we developed a method similar to those commonly used with matrix decompositions.^64^ The method works by unfolding the starting tensor along a given mode and computing the SVD to an increasing number of factors. This test then evaluates the increase in explained variance when adding the N^th^+1 factor and compares this to the increase in variance observed in a randomly permuted dataset. When adding an additional factor no longer explains more variance than by random chance, then it is assumed that additional factors represent random noise. A p-value can also be calculated by running the permutation procedure many times to generate a null distribution for comparison. The permutations are done by randomly reassigning cells to donors before tensor formation.

### Procedure for stability analysis

To test the stability of our decomposition, we designed a procedure whereby the data tensor is subsampled to some fraction of donors. For our adult SLE analysis, we subsampled to 85% of the donors. We then recomputed the decomposition to the same number of factors. The new factors found on the subsampled data were then linked back to the original factors by identifying, for each original factor, which new factor had the highest absolute value correlation with it. We repeated this procedure 500 times and report the max correlations for each original factor with the new factors from the subsampled data.

### Procedure for GSEA

To compute enriched gene sets among genes prioritized within each cell type for a given factor, we use the R package FGSEA.^65^ In tests of applying GSEA directly to an individual column of a factor loadings matrix, we noticed some spurious results appear as a result of the low-magnitude non-significant loadings being biased toward either positive or negative loadings. To avoid getting these false-positive hits, we compute a new value for each gene. This is calculated as the sum of unit scaled expression values multiplied by donor scores as follows:

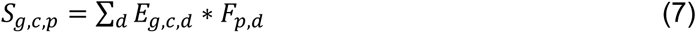

Here, *S_g,c,f_* is the new score for gene *g* in cell type *c* for factor *p*. *E_g,c,d_* is the scaled expression of gene *g* in cell type *c* for donor *d*. And *F_p,d_* is the donor score for factor *f* and donor *d*. We then apply GSEA to the new gene scores for each cell-type factor combination and apply BH multiple hypothesis test corrections. This has shown to be much more robust in preventing false-positive enriched gene sets, while still yielding gene sets that are expected based on the significant genes.

We also use GSEA when comparing similar factors across datasets as done in Figures S5C, S5D, S5E, and 5F. Before running GSEA, we first computed linear model coefficients for predicting expression from donor scores for a given factor of interest. Before computing the model, we scaled donor scores and expression across donors to unit variance and applied mean centering on 0. We only tested genes that were expressed in at least 5% of donors for a given cell type in both datasets. Next, we computed the difference in betas for each gene across datasets and squared them. This would make genes with similar factor associations across datasets have values near 0 and genes with different effects across datasets have large positive values. Then, we ran GSEA on these scores using the Cacoa package, which also clusters the gene sets in two stages for visualization^66^. The first stage clusters gene sets by similarity in the genes they contain. Secondly, these meta-sets are clustered again based on their similar patterns of enrichment across the different cell types. The number of meta-sets clustered together is shown on the plot as a prefix to the description of the most common GO terms in the set.

### Procedure for cell proportion analyses

To systematically identify shifts in subtype composition, we used either existing subtype annotations or tested various subclustering resolutions per major cell type. For the latter, we first aligned cells across the 10X lane batch variable using Conos.^7^ Then we subclustered each major cell population to varying resolutions using the *findSubcommunities* function from Conos. To get marker genes that distinguish the cell subclusters, we used the *getDifferentialGenes* function. Marker genes were plotted using the *DotPlot* function in Seurat.^67^ To test for associations between a factor and cell subtype composition, we calculated the cell proportions for each donor as the number of their cells from a given subcluster divided by the total number of cells from the corresponding major cell cluster. Next, we convert these proportions to balances using the isometric log-ratio transformation.^68^ This converts the dependent set of proportions to an independent set of p-1 variables, where p is the number of cell subtypes for a given major cell cluster. By making this conversion, it allows one to use these variables with standard statistical tests that require covariates to be independent. As a note, we also add a pseudocount of 1 to each cell proportion numerator to avoid infinities when calculating balances. Then, we use the balances as explanatory variables in a multiple linear regression against the donor scores of each factor. An F-test is used to determine whether a given cell type composition is significantly associated with a factor. The procedure is the same for computing the significance of factor associations with the overall major cell type composition.

### Procedure for LR analysis

For this analysis, we used a database of protein ligands and receptors from CellChat.^69^ We first identified clusters of co-expressed genes in the pseudobulk data for each cell type using the R package WGCNA.^20^ Specifically, we used the signed network and TOM similarity matrix with a module tree cut height of 0.25. Then, we calculated the association between each module eigengene and each ligand from the list of cognate LR pairs. This is only done if the receptor was present in the module cell type to some minimal level (expressed in at least 5 cells for each of the top 15% of donors by ligand expression). Filtering was applied such that we only tested ligands where at least 1% of donors have scaled-adjusted expression above 0.2 (the same is applied to both ligands and receptors in the LR association test below). This removed ligands where high expression is observed in only a few donors or if the ligand has low normalized variance across donors. We also only tested for interactions between different cell types, as we would likely find many false-positive associations between ligands and modules found in the same cell type (due to regulation of the ligand by the same upstream transcription factor regulating the module). The association p-values were calculated by a linear model F-tests with the module eigengene as the response variable and the scaled expression of a ligand as the explanatory variable. P-values were adjusted using the BH procedure. When the CellChat database listed multiple receptor components as required for a specific LR interaction, we required all components of the receptor complex to be expressed. The heatmap in Figure 4B only includes rows with at least one adjusted p-value below 5.0×10^-11^ and columns with at least one adjusted p-value below 0.001 to reduce the number of results for visual purposes. Finally, we computed the overrepresentation of gene sets within gene co-expression modules of interest. We employed the hypergeometric test to determine whether a module contained a significantly higher proportion of genes from specific gene sets compared to the rest of the genes not found in the module but still included in the analysis. For the B_m1 enrichment tests, we included gene sets from GOBP, KEGG, and Reactome databases. For benchmarking our approach, we compared it to a standard LR association method and a random selection of LR channels. The LR association test inferred an LR interaction if the expression of the ligand from one cell type was significantly associated (adjusted p-value < 0.05) with the expression of its cognate receptor in another cell type. For the scITD LR inference method, we used a more stringent adjusted p-value of 0.0001 to make the number of inferred interactions roughly the same for at least one of the LR pair databases (CellChat database in this case).

To demonstrate that our method could enrich for a more confident set of LR interactions, we used NicheNet regulatory potential scores for each ligand. For a given LR channel consisting of a ligand, a source cell type, and a target cell type, we first identified the top 200 genes in the target cell type by the highest absolute value Pearson correlation with the ligand’s expression in the source cell type. Then, we compared the regulatory potential scores for these top 200 genes with those from all other genes. We used a Wilcoxon rank-sum test to test for differences in regulatory potential scores between these two groups. As a note, we also applied the *normalize_correlation* function from the *spqn* package to correct for gene-gene correlation biases with mean expression.^70^

We also directly compared the performance of our LR analysis method with that of the Differential NicheNet method, which is also designed to determine differentially active LR interactions across samples. To assess performance, we relied on data from 101 publicly available ligand treatment datasets for 47 unique ligands, which were also used as benchmarking in the NicheNet study. In this pipeline, we first ran scITD to infer differentially active LR interactions (Padj < 0.05) across donors in factors aSLE_F1, aSLE_F2, and aSLE_F3. Then, we ran Differential NicheNet for the same factors, selecting the same number of top results. As this version of NicheNet only runs on data in the case-control scenario, we needed to first discretize our factors prior to running the tool. To discretize factors, we selected the top and bottom quintile of donors by donor scores as the case and control groups. For a given interaction channel involving a ligand and target cell type, we aimed to test whether the factor associated genes in the target cell type were enriched for genes that became differentially expressed in one of the corresponding ligand treatment datasets. Therefore, we condensed our results from each method to only those interaction channels where the ligand was present in one of the ligand treatment datasets. For each of these channels, we selected the significantly differentially expressed genes from the corresponding ligand treatment dataset as genes with both absolute value log2FC > 1 and qvalue <0.1. Using these genes as a gene set, we tested whether the factor-associated genes were enriched for these genes. This was computed using FGSEA applied to the square of linear model coefficients for all genes above a baseline expression threshold with the respective factor. In cases where multiple ligand treatment datasets were available for the same ligand, we selected the enrichment with the best p-value and applied a Bonferroni correction for the number of datasets tested. After all enrichment tests were calculated we applied an FDR correction to all results generated from each method. We then computed the fraction of NicheNet results or scITD results that had a significant enrichment (Padj<0.001). For each method, we also selected 10 random sets of channels from the ligands in the ligand treatment datasets. These random sets were constructed to be the same size as the number of channels tested for each method, so that the fraction and variance of significant results could be directly compared to the real results.

### Nehar-Belaid et al. pediatric SLE data preprocessing and analysis

We preprocessed the data the same way as in the original study.^30^ We combined fine-resolution cluster assignments to match the cell type resolution analyzed in the other datasets that we analyzed. We determined the appropriate number of factors for the decomposition using our stability analysis described above. To identify factors from this dataset corresponding with aSLE factors (Figure S2H), we projected this data onto the aSLE factors and calculated pearson correlations between the donor scores from the projection and those from the pSLE decomosition. To compare the loadings between cognate pediatric and adult factors (Figure S5A), we first computed the linear model coefficients for all genes associations each factor (only for genes expressed in at least 5% of donors). Then, for a given factor comparison, we selected the union of genes that were significant in either dataset (Padj<.05) and computed the Pearson correlation of linear model coefficients across genes. The clinical association tests were carried out the same as in the adult SLE dataset depending on the variable type. In total, we tested 10 metadata associations with specific projected factor scores determined by our adult SLE metadata associations. The meta-analysis p-values were determined by Fisher’s method combining each nominal pediatric association p-value with its corresponding nominal p-value from the adult SLE association tests and applying FDR correction afterwards assuming 10 tests.

### Stephenson et al. COVID-19 data preprocessing and analysis

We used the quality-controlled data that was made publicly available.^46^ A few donors had multiple samples taken at various time points. We kept only the samples labeled with collection day “D0”, so that the dataset contained 1 sample per donor. We also excluded patients with other diseases besides COVID-19, and we also excluded the healthy donors given LPS. We used the previous annotations labeled as “full_clustering” and grouped cell subtypes to form the major cell type clusters. Specifically, for B cells we included “B_exhausted”, “B_immature”, “B_naive”, “B_non-switched_memory”, and “B_switched_memory”. For cMonocytes, we included “CD14_mono” and “CD83_CD14_mono”. For Th cells we included “CD4.CM”, “CD4.EM”, “CD4.IL22”, “CD4.Naive”, “CD4.prolif”, “CD4.Tfh”, and “CD4.Th1”. For Tc cells, we included “CD8.EM”, “CD8.Naive”, “CD8.Prolif”, and “CD8.TE”. For NK cells we included “NK_16hi”, “NK_56hi”, and “NK_prolif”. We also noticed that some of the cells previously labeled as “B cells” clustered with the plasmablasts and expressed the same gene and protein markers as the plasmablasts. Therefore, these were excluded from the B-cell pseudobulks. After removing donors with less than 2 cells in any of the included cell types, we were left with 103 donors and 452,740 cells from the major cell populations used in the analysis. The median number of cells per donor for B, Tc, Th, NK, and cMonocytes were 386, 558, 573, 593, and 717 cells, respectively. The samples were processed at one of three different sites including Cambridge, NCL, and Sanger. Therefore, we applied ComBat batch correction to account for these technical differences as was done with the SLE dataset. We ran the Tucker decomposition to 9 donor factors and 26 gene factors using our hybrid rotation method. We further used the FGSEA package for computing enrichment of donors by status (COVID-19, healthy, or COVID-19 critical) in either high or low CVD_F1 scores. For testing enrichment of critical patients, we excluded the healthy donors. We compared these factors to those from the adult SLE dataset using the same approach outlined above to compare the pediatric and adult SLE datasets. To check that dataset differences were not due to age- and sex-imbalance between datasets, we downsampled the COVID-19 and SLE datasets to match the proportions of these donor metadata. To achieve roughly the same representation, we first randomly subsampled the COVID-19 dataset to keep only 10% of male donors. Then, we randomly subsampled donors from each age bin of each dataset to roughly match age proportions.

### van der Wijst et al. COVID-19 data preprocessing and analysis

We used the quality-controlled data that was made publicly available.^16^ We removed cases that were negative for SARS-CoV-2. We only used samples measured at day 0, such that we retained 1 sample per donor. Broad cell clusters and annotations were used from the original study. After removing donors with less than 2 cells in any of the included cell types, we were left with 60 donors and 166,970 cells from the major cell populations used in the analysis. The median number of cells per donor for B, Tc, Th, NK, and cMonocytes were 208, 459.5, 637, 278.5, and 747 cells, respectively. We also applied ComBat batch correction at the level of 10X lanes. Finally, we ran the Tucker decomposition to 10 donor factors and 30 gene factors using our hybrid rotation method. We projected this data onto the CVD_F2 pattern from the Stephenson et al. dataset the same way as the other projections (described above). We used Fisher’s method to compute the meta-analysis p-value combining the severity-association test with that of the larger dataset.

### Practical use tips for scITD R package

The minimum required input to run scITD is a gene-by-cells UMI counts matrix and metadata matrix indicating the cell type and source donor of each cell (dimensions of cells-by-2 columns). Additional columns in the latter matrix may be included to test for associations between the factors and donor-level variables. To generate the cell type annotations, we recommend using one of the many sample-alignment methods currently available to establish correspondence between cell types across samples prior to clustering and identifying marker genes. However, it is crucial that the input UMI matrix uses the original, non-sample-aligned data, otherwise sample-derived expression variation may be removed. The UMI counts matrix should also be cleaned of potentially empty droplets, doublets, or of cells with a high percent of UMIs from mitochondrial genes before being input to scITD.

Furthermore, it is optimal (but not required) for the data to be generated by pooling and demultiplexing,^3^ as this enables more effective elimination of batch effects by removing confounding between sample and batch. Our tool will apply batch effect correction automatically using ComBat if the user specifies a batch covariate. It is also necessary to specify other preprocessing parameters in *form_tensor()*, including a minimum number of cells per donor, the type of normalization to use (standard or EdgeR trim mean normalization), a normalization scale-factor, a cutoff for selecting top overdispersed genes, and a variance scaling parameter. In the package, we provide default values that will work well for most scenarios.

For running the tensor decomposition, we provide a function, *run_tucker_ica().* The user needs to specify the number of factors to extract as well as the type of rotation to apply (ICA on donor scores or hybrid rotation on loadings). We also provide a function, *determine_ranks_tucker()*, to help the user determine an appropriate number of factors to use in *run_tucker_ica()*. This implements the SVD-based significance test we described in the text. Other functions are included to help evaluate the quality of decomposition including *run_stability_analysis()* for evaluating stability of the decomposition to downsampling donors. The user can choose to compute linear associations between donor scores and metadata with *get_meta_associations()*. To visualize factors, we have a function to create a heatmap of the donor scores matrix called *plot_donor_matrix()* and a function called *plot_loadings_annot()* to create a heatmap the loadings matrix for a given factor. These visualizations and many others in the package are generated using the ComplexHeatmap package.^71^

For post processing the factors with GSEA, we include the function *run_gsea_one_factor()*. To run the LR analysis, we require input of a matrix with two columns specifying cognate ligand-receptor pairs. We include functions to prepare the data for this analysis, generate the WGCNA gene modules, and to run the analysis called *prep_LR_interact()*, *get_gene_modules()*, and *compute_LR_interact()*, respectively. Both the GSEA and LR functions output heatmap plots of the results.

## Data Availability

The IFN-beta stimulation dataset can be found in the Gene Expression Omnibus (GEO) at accession number GSE96583. The count-matrices for the Perez et al. SLE scRNA-seq dataset are available in GEO at accession GSE137029. The Nehar-Belaid et al. pediatric SLE scRNA dataset was downloaded from GEO at accession GSE135779. The Stephenson et al. COVID-19 dataset is publicly available at https://www.covid19cellatlas.org/index.patient.html, titled “COVID-19 PBMC Ncl-Cambridge-UCL”. The van der Wijst et al. COVID-19 dataset can be found at https://cellxgene.cziscience.com/collections/7d7cabfd-1d1f-40af-96b7-26a0825a306d.

## Code Availability

Our computational method, scITD, can be found at https://github.com/kharchenkolab/scITD <u>or</u> on The Comprehensive R Archive Network (CRAN) at https://cloud.r-project.org/web/packages/scITD/index.html. A walkthrough tutorial for using scITD is also available at http://pklab.med.harvard.edu/jonathan/. The code used to produce all figures in this paper can be found at https://github.com/j-mitchel/scITD-Analysis/tree/main/figure_generation.

## Author Contributions

P.V.K. has formulated the study and with J.M. developed the overall approach. J.M. developed the detailed algorithms with advice from P.V.K. with assistance from E.B. J.M., C.J.Y., and P.V.K. worked on interpretation of results, with help from M.G.G., R.K.P., and R.B. J.M. and P.V.K. drafted the manuscript, with contributions from C.J.Y. and input from other authors.

## Competing Interests

P.V.K. serves on the Scientific Advisory Board to Celsius Therapeutics Inc. and Biomage Inc. P.V.K. is an employee of Altos Labs. C.J.Y. is a SAB member for and hold equity in Related Sciences and ImmunAI, a consultant for and hold equity in Maze Therapeutics and a consultant for Trex Bio. C.J.Y. has received research support from Chan Zuckerberg Initiative, Chan Zuckerberg Biohub and Genentech.

## Acknowledgements

We thank Anna Igolkina (St. Petersburg Technical University, Russia) for advice on the cell proportion analysis.

---

**Figure S1:**
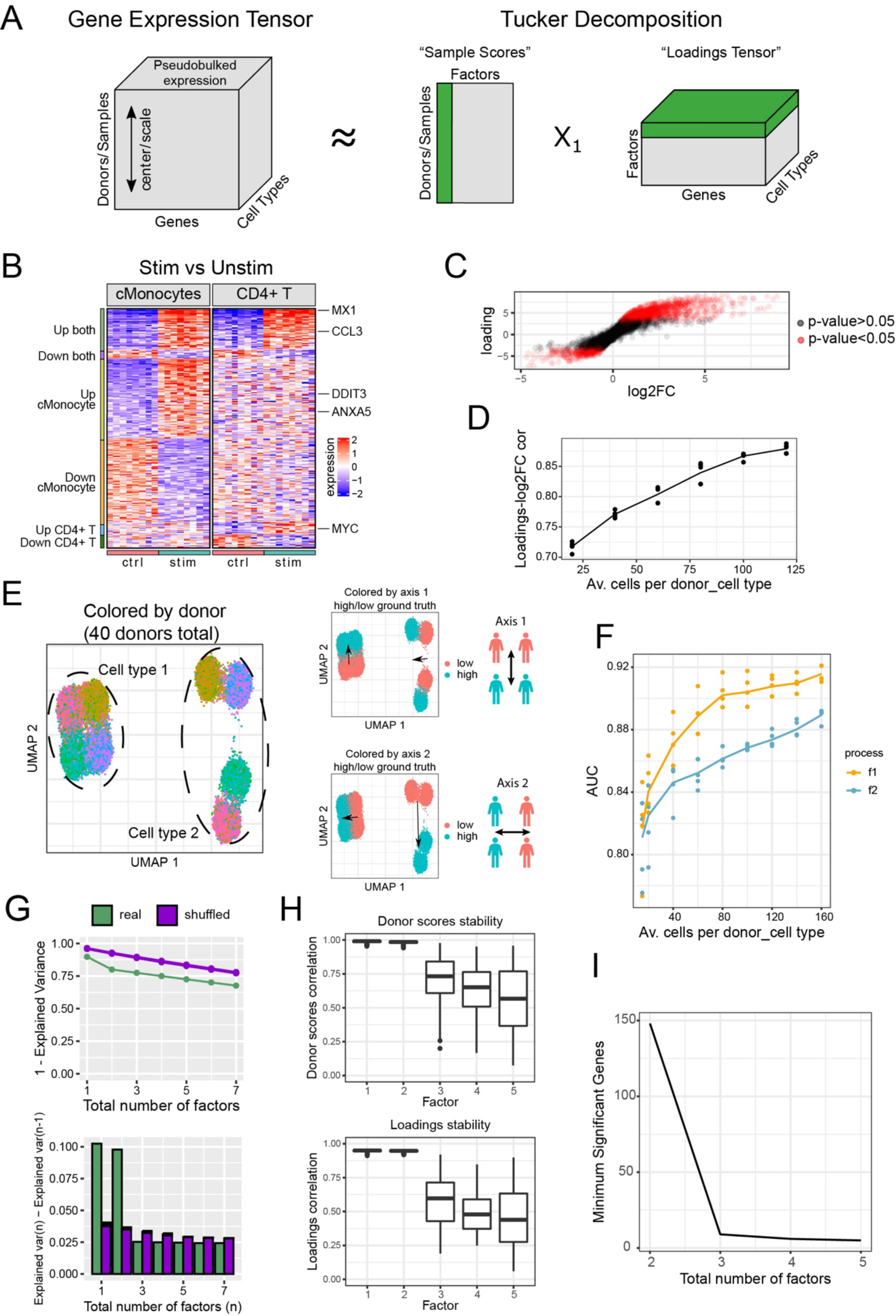
scITD decomposition overview and additional validation results. (A) Structure of the output from scITD (middle and right) applied to a single-cell pseudobulk expression tensor (left). An approximation of the expression tensor is reconstructed when the sample scores matrix (middle) and loadings tensor (right) are multiplied together. Sample scores and loadings for one factor are highlighted in green. (B) Sample-level pseudobulk gene expression of DE genes between control and IFN-beta stimulated samples. Rows are genes and columns are pseudobulked samples. Genes that are significant in at least one of the two cell types below an adjusted p-value of 0.01 were included. Genes are grouped (left annotation) by the cell types where they were DE across conditions. A few DE genes are shown labeled on the right. (C) Comparison of DE log2 fold-change values (from D) to loadings values from factor 1. Each point is a gene in one of the two cell types. The Spearman correlation is shown in the upper-right corner. Red dots represent genes that are significantly associated with factor 1 at an adjusted p-value < 0.05. (D) Spearman correlations between loadings and log2FC from the IFN-beta experiment data downsampled to a varying average number of cells per donor per cell type. Each point represents a different downsampling iteration (n=5), and the line is the mean Spearman correlation at a given dataset size. (E) UMAP of the simulated scRNA-seq dataset with two cell types (left). Also shown are two multicellular patterns that separate groups of donors (right). Arrows point to the cells from donors with upregulation of a given multicellular pattern. (F) Performance of the method on the simulated dataset downsampled to a varying average number of cells per donor per cell type. AUC is calculated from predicting ground truth DE genes in each cell type from each gene’s expression-factor association p-value. AUC is shown for each multicellular pattern as distinct colors. Each point represents a different downsampling iteration (n=5), and the line is the mean AUC at a given dataset size. (G) Rank determination by SVD applied to the full simulated dataset (Methods). The top plot shows the total explained variance percent when applying SVD to varying numbers of factors. The bottom plot shows the change in explained variance percent when incrementing the total number of factors. Green bars show the results for the simulated dataset, whereas purple bars show the results for the dataset after randomly shuffling cell- to-donor assignments (n=50 shuffling iterations). Error bars for the shuffled samples represent standard deviation. (H) Stability analysis results for the simulated data decomposition over 500 iterations. In each iteration, the dataset was subsampled to 70% of the donors. Values represent the maximum absolute value factor-factor correlation coefficients for each of the original factors mapped to the factors from each subsampled dataset decomposition. This is shown separately for max donor score correlations (left) and max loadings correlations (right). (I) The y-axis indicates the minimum number of significant genes per-factor from factors of a given decomposition. This is shown for decompositions to differing numbers of total factors.

**Figure S2.**
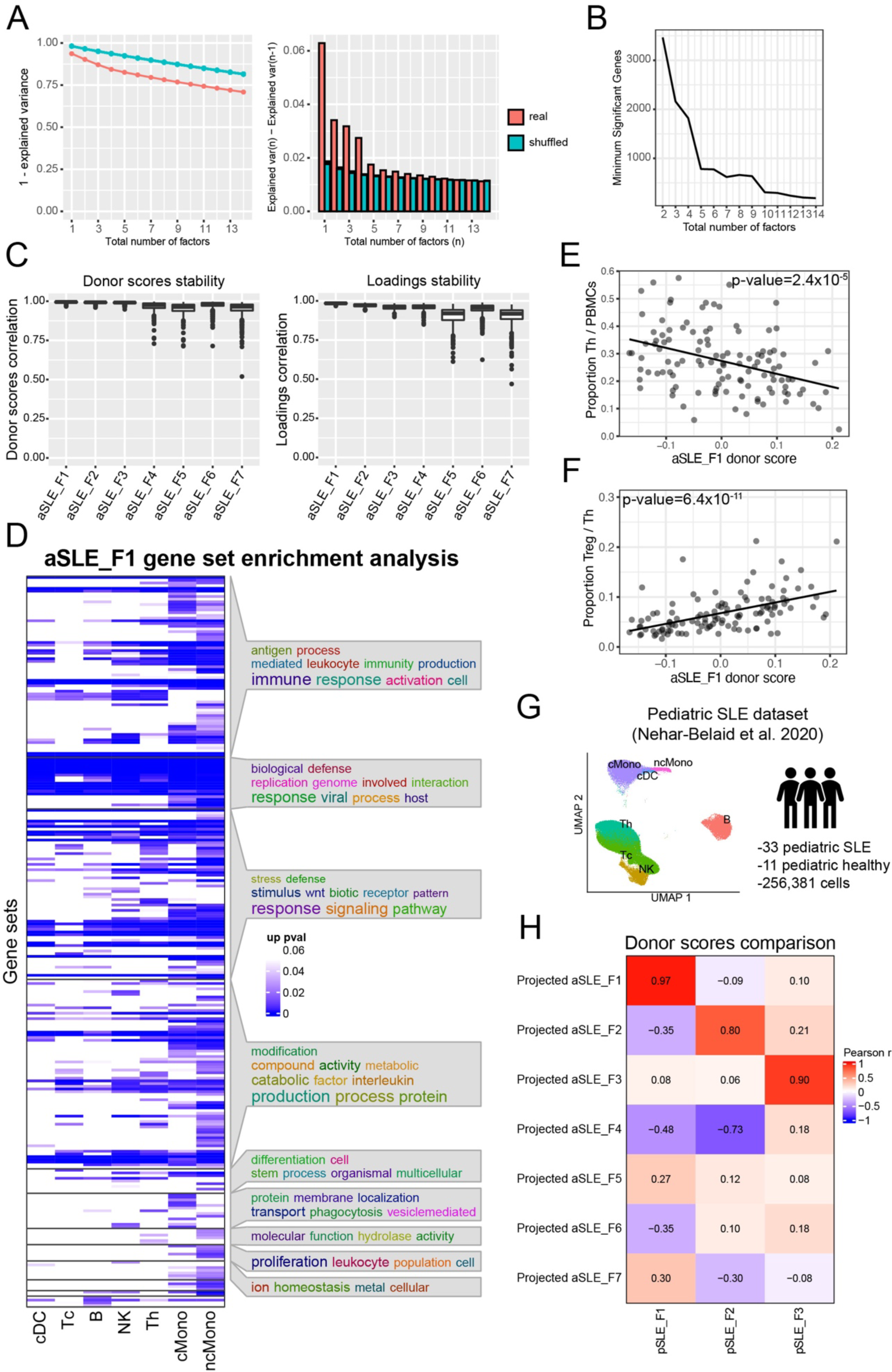
Additional results for aSLE_F1 and overall factor similarity to pediatric dataset decomposition. (A) Rank determination by SVD applied to the adult SLE dataset (Methods). The top plot shows the total explained variance percent when applying SVD to varying numbers of factors. The bottom plot shows the change in explained variance percent when incrementing the total number of factors. Red bars show the results for the real dataset, whereas blue bars show the results for the dataset after randomly shuffling cell-to-donor assignments (n=50 shuffling iterations). Error bars for the shuffled samples represent standard deviation. (B) Minimum number of significant genes among all factors, for incrementally increasing total numbers of factors in the decomposition. A linear model F test is used to determine the number of genes associated with each factor. (C) Stability analysis results for the SLE-only decomposition over 500 iterations. In each iteration, the dataset was subsampled to 85% of the donors. Values represent the maximum absolute value factor-factor correlation coefficients for each of the original factors mapped to the factors from each subsampled dataset decomposition. This is shown separately for max donor score correlations (left) and max loadings correlations (right). (D) Gene set enrichment for aSLE_F1 factor, computed for each cell type separately (Methods). Gene sets clustering and word clouds generation were implemented using the simplifyEnrichment R package. (E) Association of the total fraction of Th cells with aSLE_F1 scores. The line is a linear model and significance was calculated by the linear-model F-test (also applies to D). (F) Association of the fraction of Treg cells out of all Th cells with aSLE_F1 scores. (G) Description of pediatric SLE dataset and UMAP of cell clusters used. (H) Correlations between projected donor scores (using aSLE factors) and pediatric donor scores.

**Figure S3.**
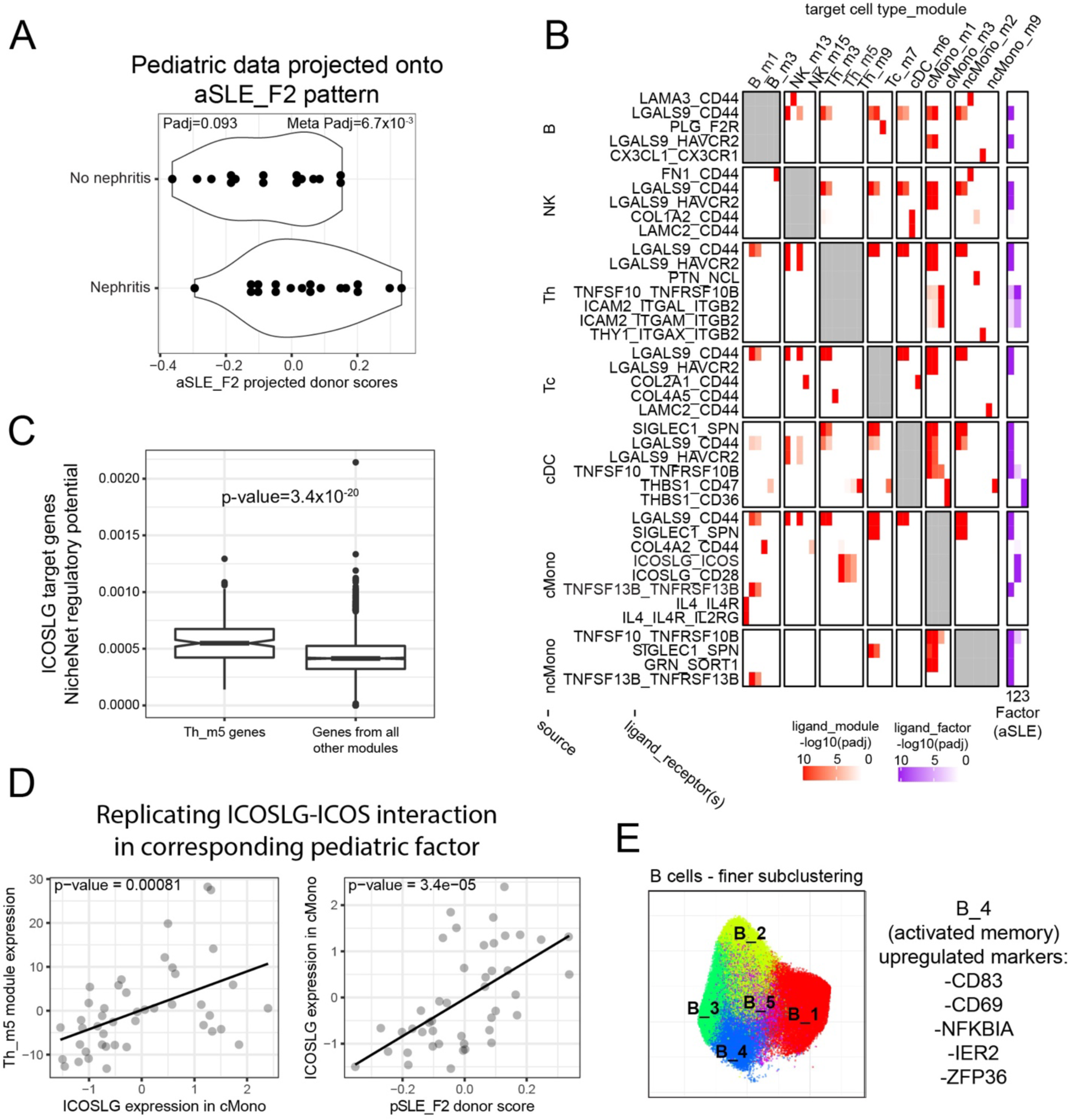
Additional results for the nephritis-associated factor and ligand-receptor interaction inference. (A) Replicating the nephritis associated signal in the pediatric dataset. The pediatric data were projected onto the aSLE_F2 multicellular pattern, and the resulting donor scores were plotted against nephritis metadata. Logistic regression with a likelihood-ratio test was used to evaluate significance. (B) Additional top results of the scITD LR analysis using the CellChat LR pair database (left). Rows are ligand hits from various source cell types. Columns are gene co-expression modules from various target cell types. Rows are grouped by source cell type and columns are grouped by target cell type. Values in the main body of the heatmap indicate adjusted p-values for ligand-module associations. Only the top significant results are shown (Methods). Rows and columns are clustered within each block. Also shown are ligand-factor association adjusted p-values (right). (C) ICOSLG-target gene regulatory potential scores (from NicheNet) are shown for genes in the Th_m5 module or all other modules. The Wilcoxon rank-sum test was used to test for a difference in medians between the two groups. (D) Replicating the ICOSLG LR interaction in the pediatric dataset with the corresponding pediatric factor, pSLE_F2. The same genes from the Th_m5 of the adult SLE dataset were used to compute gene module scores. (E) Finer grained subclustering of B cells in UMAP space and marker genes distinguishing B_4 as an activated memory subtype.

**Figure S4.**
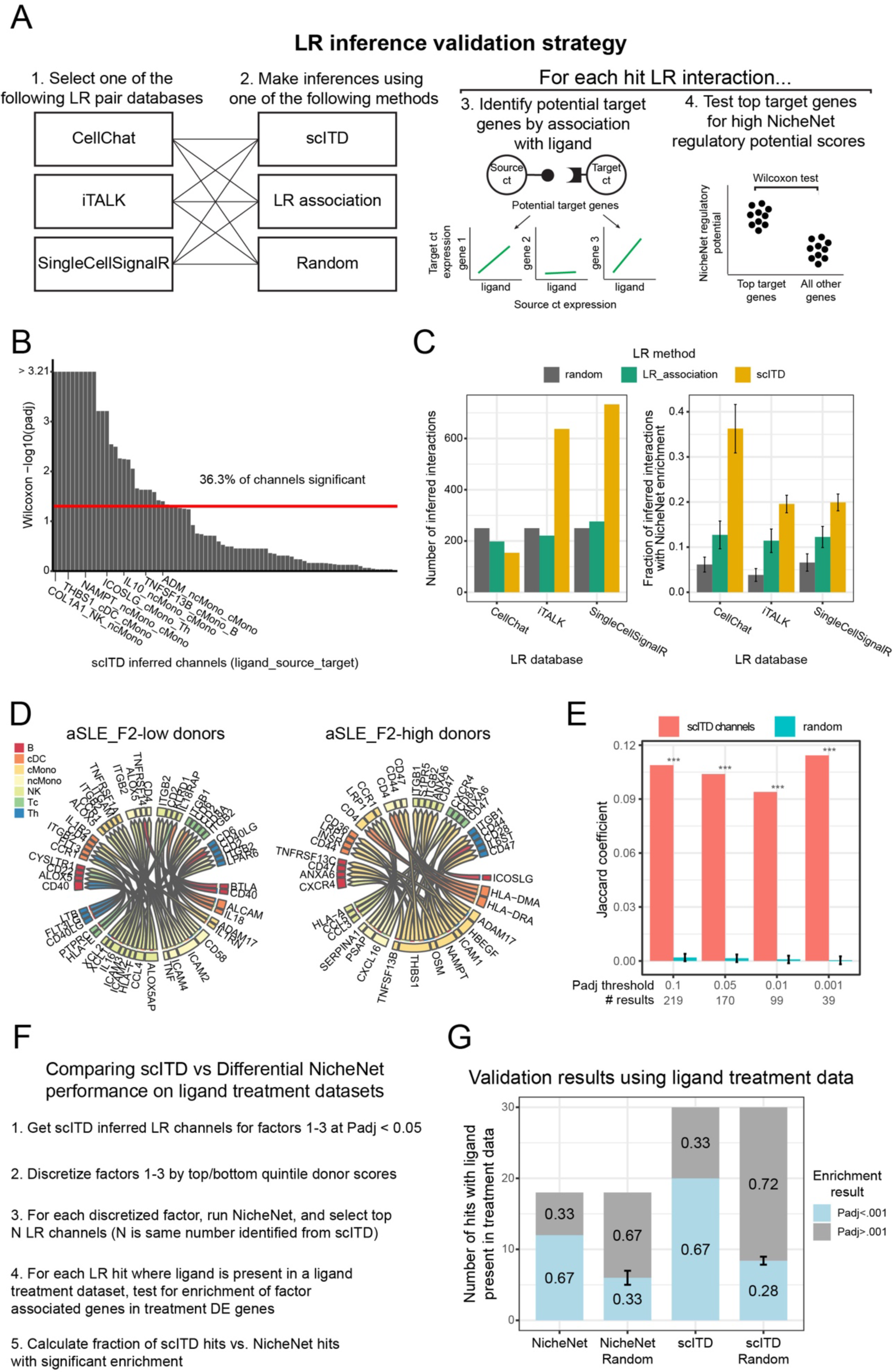
Validation of the LR inference approach using NicheNet and comparison of scITD results to Differential NicheNet. (A) Overview diagram of the strategy used to validate our LR interaction inference approach using external data from NicheNet. (B) LR channels inferred using scITD with the CellChat LR pair database. The height of each bar indicates the adjusted p-value (calculated with the Wilcoxon rank-sum test) comparing NicheNet regulatory potential for the top 200 potential target genes to the rest (Methods). The red bar indicates an adjusted p-value of 0.05. (C) The number of inferred LR channels using scITD compared to a simple LR pair association approach (left). The plot on the right shows the fraction of inferred LR channels with significant NicheNet enrichment (as calculated in G). The “random” method is a random selection of 250 LR channels as a background comparison. Error bars are the standard error of the mean. (D) Top results from Differential NicheNet analysis. The tool was run on the top and bottom quintile donors (by donor score) stratified by aSLE_F2. (E) Overlap between Differential NicheNet results and scITD results associated with aSLE_F2. Significance was evaluated by a Z-test compared to the null distribution. (F) Overview of analysis pipeline used for comparing performance of scITD and NicheNet using ligand treatment datasets. (G) Comparison betwee scITD and NicheNet, showing the fraction of hits with supporting evidence from the ligand treatment datasets. “NicheNet Random” and “scITD Random” refer to a random selection of LR channels, matching the same number of tests from either method. Error bars represent standard error for the fraction of random tests below the p-value threshold.

**Figure S5.**
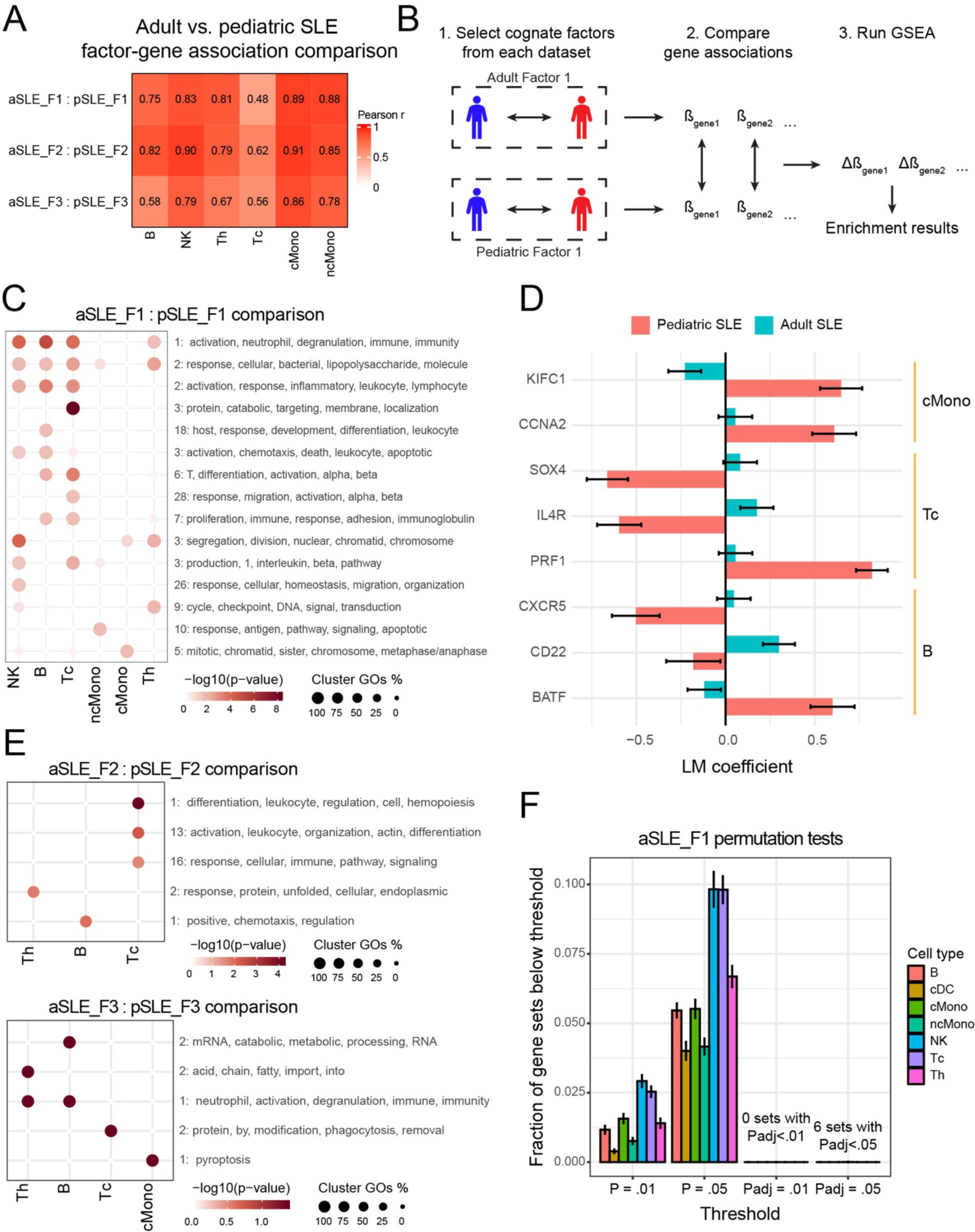
Differences between the corresponding adult and pediatric SLE factors. (A) Pearson correlations of gene-factor association coefficients between aSLE and pSLE datasets. Only genes that had significant associations with a given factor in either dataset were used for computing the correlations. (B) Schematic diagram illustrating the general approach used to compare similar factors (Methods). (C) Unsigned gene set enrichment analysis using dataset-specific upregulated or downregulated genes when comparing aSLE_F1 versus pSLE_F1. Gene sets were collapsed in two stages, first by overlapping sets of genes and then by the pattern of gene sets across cell types. The number in front of the gene set terms indicates the number of sets of gene sets with similar terms that clustered together (Methods). (D) Factor-gene expression association linear model coefficients for select leading edge genes from significant gene sets in (C). (E) Unsigned gene set enrichment analysis using dataset-specific upregulated or downregulated genes when comparing aSLE_F2 versus pSLE_F2 (top) or when comparing aSLE_F3 versus pSLE_F3 (bottom) as in (C). (F) Negative control for the factor comparison gene set enrichment tests. We subsampled the aSLE dataset to 90% of donors, reprocessed the data, reran scITD, and followed the procedure in (B). This was repeated 30 times. Bar heights represent the mean fraction of gene sets with a p-value below each threshold, and error bars represent standard deviation.

**Figure S6.**
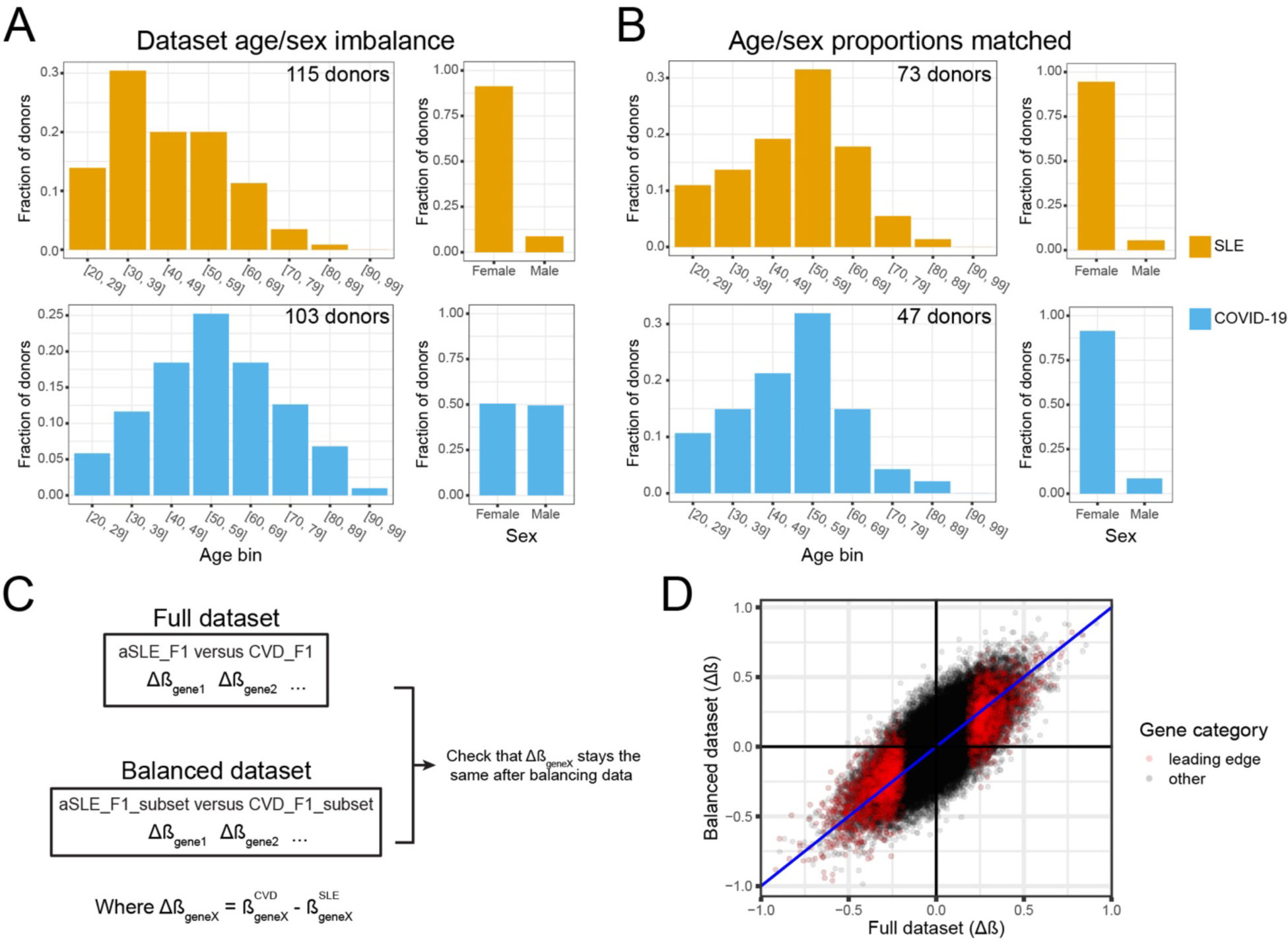
Evaluating impact of dataset imbalance on SLE versus COVID-19 comparison. (A) Age and sex distribution for donors of the SLE dataset (top) versus COVID-19 dataset (bottom). (B) Age and sex distribution for donors of the SLE dataset (top) versus COVID-19 dataset (bottom) after downsampling the datasets to match these characteristics (Methods). (C) Approach taken to evaluate the impact of dataset imbalance on the analysis of gene-association statistic differences across datasets (Methods). (D) Results from the strategy outlined in (C), showing the difference in association statistics from the original comparison versus after balancing the datasets for age and sex. Each point is a gene tested in a cell type, with the red dots indicating those genes that contributed to at least one of the enriched gene sets in Figure 5F for the respective cell type.

**Figure S7.**
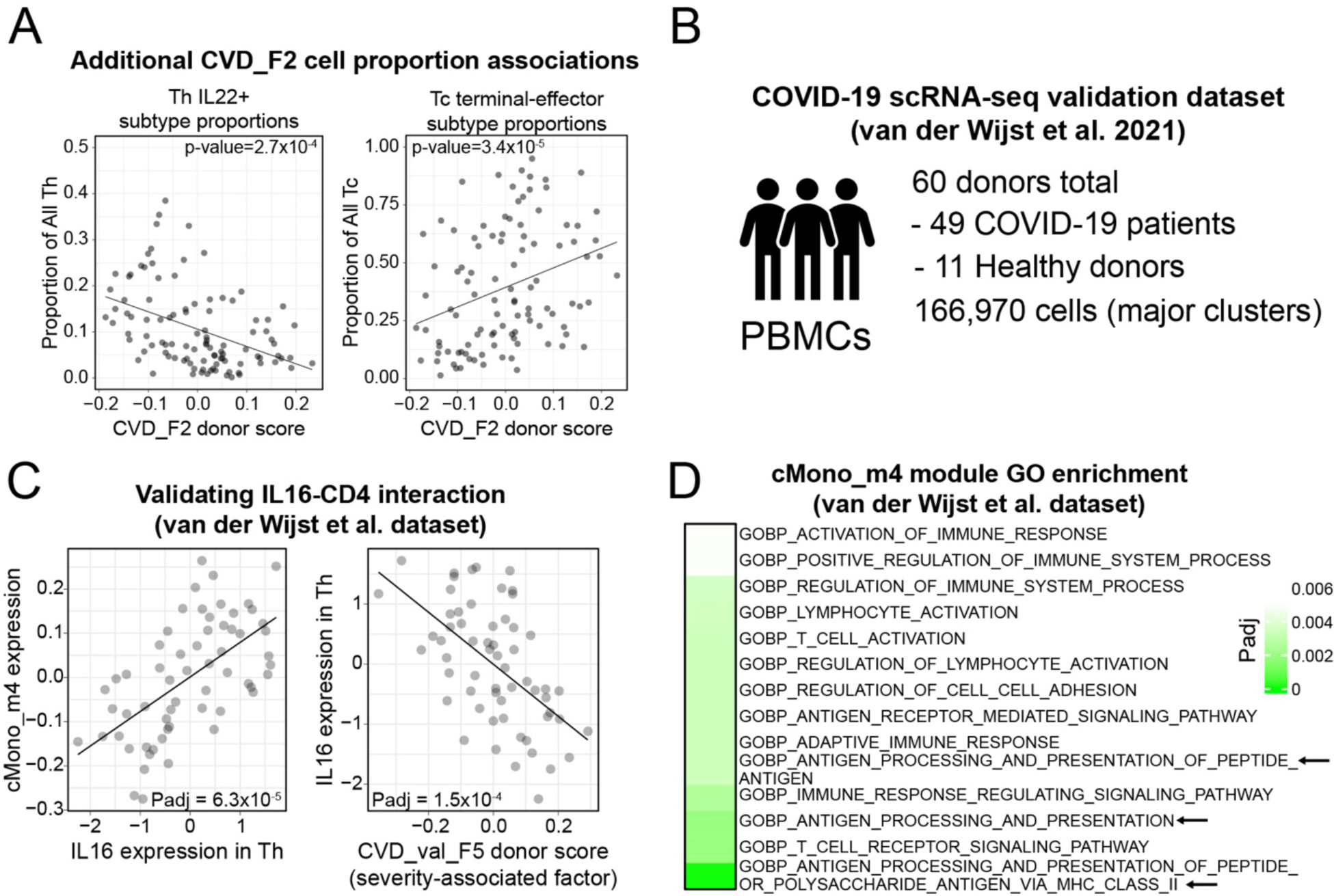
Additional COVID-19 severity associated factor results. (A) CVD_F2 associations with cell subtype proportions for IL22+ Th cells (left) and terminal effector Tc cells (right). (B) Overview of van der Wijst et al. dataset used for validating findings in the main dataset. (C) The IL16-CD4 interaction identified in the van der Wijst et al. dataset. Shown is the association between *IL16* expression in Th cells with donor scores for factor 5 (the COVID-19 severity-associated factor for this dataset) (left). Also shown is the association between gene module cMono_m4 and expression of ligand *IL16* in Th cells (right). The line is a linear model. (D) Top enriched GO gene sets in co-expression module cMono_m4 from the van der Wijst et al. dataset (adjusted p-values < 0.005).

## Supplemental information

Note S1, related to methods discussion of factor rotations and their implications

One unique attribute of the Tucker tensor decomposition is that different components of the output can be rotated to improve the overall interpretability (Unkel et al., 2011; Zhou and Cichocki, 2012). Here, we tested two different rotation approaches. The first of these is our hybrid rotation method applied to the loadings, with a counter rotation on the donor scores (Methods) (Figure A top). We will refer to this as the *loadings rotation*. The second is a rotation of ICA applied to the donor scores matrix with a counter rotation of the loadings (Figure A bottom). We refer to this as the *donor scores rotation*. In our comparisons, we apply either method to a decomposition of the full SLE dataset. For *loadings rotation* we extracted 7 factors, and for the *donor scores rotation* we extracted 9 factors because the latter yielded a more even spread of explained variance among the factors.

Firstly, we noticed that the multicellular expression patterns from the *donor scores rotation* often appeared highly similar to one another, as several had ISGs that were highly loading across all cell types. We quantified this effect by evaluating the average association (R-squared) between each factor and the expression of several core ISGs. This set of ISGs included the genes *ISG15, IFI27, IRF7, HERC5, LY6E, MX1, OAS2, OAS3, RSAD2, USP18,* and *GBP5* (Figure B). This analysis revealed ISG associations with multiple factors from the *donor scores rotation*, whereas only one factor yielded strong ISG associations in the *loadings rotation* (Figure B). We found the latter to be helpful in ascribing more succinct biological summaries to each factor (when using tools like GSEA), aiding their interpretability.

Secondly, we noticed that the *donor scores rotation* sometimes produced factors that were more strongly associated with certain metadata variables. We evaluated the association strength of the ethnicity-associated factor for the *loadings rotation* versus the *donor scores rotation* and found that the *donor scores rotation* more clearly separated the donors by ethnicity (Figure C). In summary, the *donor scores rotation* can help stratify patients if discrete patient subgroupings exist, but it can make the multicellular processes less modular and challenging to interpret compared to the *loadings rotations*.

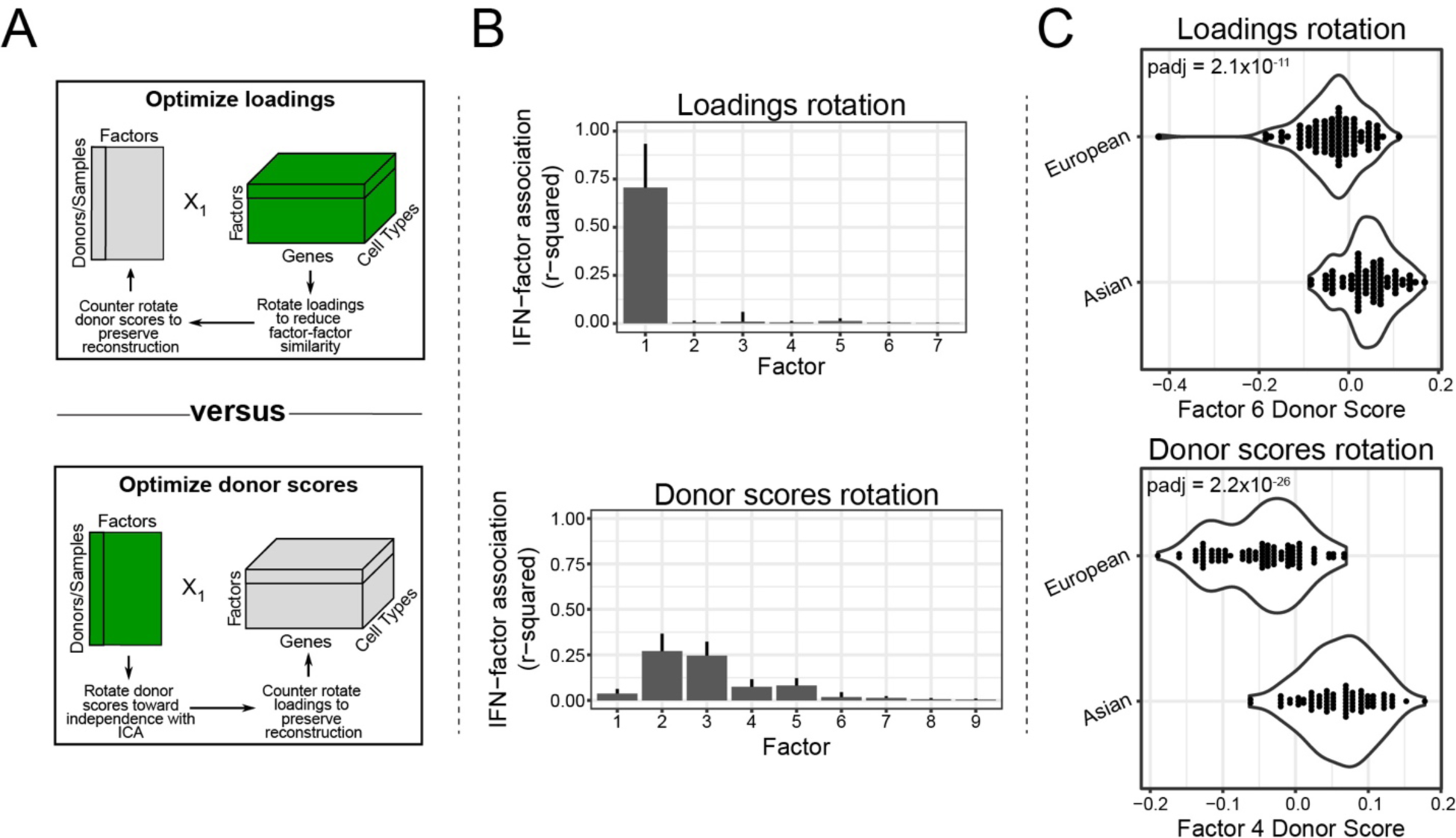
Comparing the impact of different factor rotations. (A) The two general approaches taken to rotating factors. Either the loadings are rotated to maximize some criterion (top), or the donor scores are rotated to maximize some criterion (bottom). In either case, the non-optimized component of the decomposition output is counter-rotated to preserve the original reconstruction error. (B) Average ISG-factor score associations (R-squared) for the loadings rotation (top) and donor scores rotation (bottom). The ISGs used are *ISG15, IFI27, IRF7, HERC5, LY6E, MX1, OAS2, OAS3, RSAD2, USP18,* and *GBP5*. A separate R-squared value is calculated for each cell type. The error bars represent the standard deviation across this set of genes. (C) Comparing the strength of association for the ethnicity-associated factor from the loadings rotation (top) and the donor scores rotation (bottom). The p-values were calculated using univariate linear model F-tests.

## Supplemental information

Note S2, related to discussion of alternate applications of the tool

When using scITD on the full SLE dataset without any batch correction, we were able to isolate batch effects into distinct factors (Figure A). This provides a unique opportunity to explicitly study the multicellular nature of batch effects. We further showed that 10X Chromium lane-associated factors often have expression patterns where the same genes are up/downregulated across all cell types. This is shown for Factor 1, which was associated with the batch, dmx_YE_7-19 (Figure B). This pointed us to think that this is likely due to some ambient RNA contamination in these batches. Specifically, we found that the 10X lane-associated genes shared between cell types were more likely to be present at higher fractions in droplets containing only ambient RNA (droplets with ≤ 10 UMIs) compared to the cell-type-specific genes (Figure C). This analysis provides a unique use case for scITD to examine how batch effects impact multiple cell types and may eventually pave the way for improved batch-correction techniques.

Finally, we analyzed whether the genes involved in the batch-associated factors tended to be higher or lower in GC content. GC content from each gene was retrieved using the EDASeq package in R (Risso et al., 2011). To test whether the upregulated genes of a given batch factor had a significant association with GC content, we used a two-sample t-test comparing GC contents for upregulated genes versus all other genes included in the analysis. For the non-batch factors, we arbitrarily selected the positive loading direction to call upregulated genes. Many of the batch associated factors showed strong biases for high/low GC content genes, whereas no such biases were observed in the non-batch associated factors (Figure D). After applying batch correction at the pool-level and using the same decomposition parameters, these GC content associations were greatly reduced in strength (testing for associations in both directions), though some remained statistically significant. The strongest of these post-correction GC content associations had a p-value of 0.002, with 8/25 factors having adjusted p-values between 0.01 and 0.002. However, these associations are orders of magnitude weaker compared to those from before batch correction.

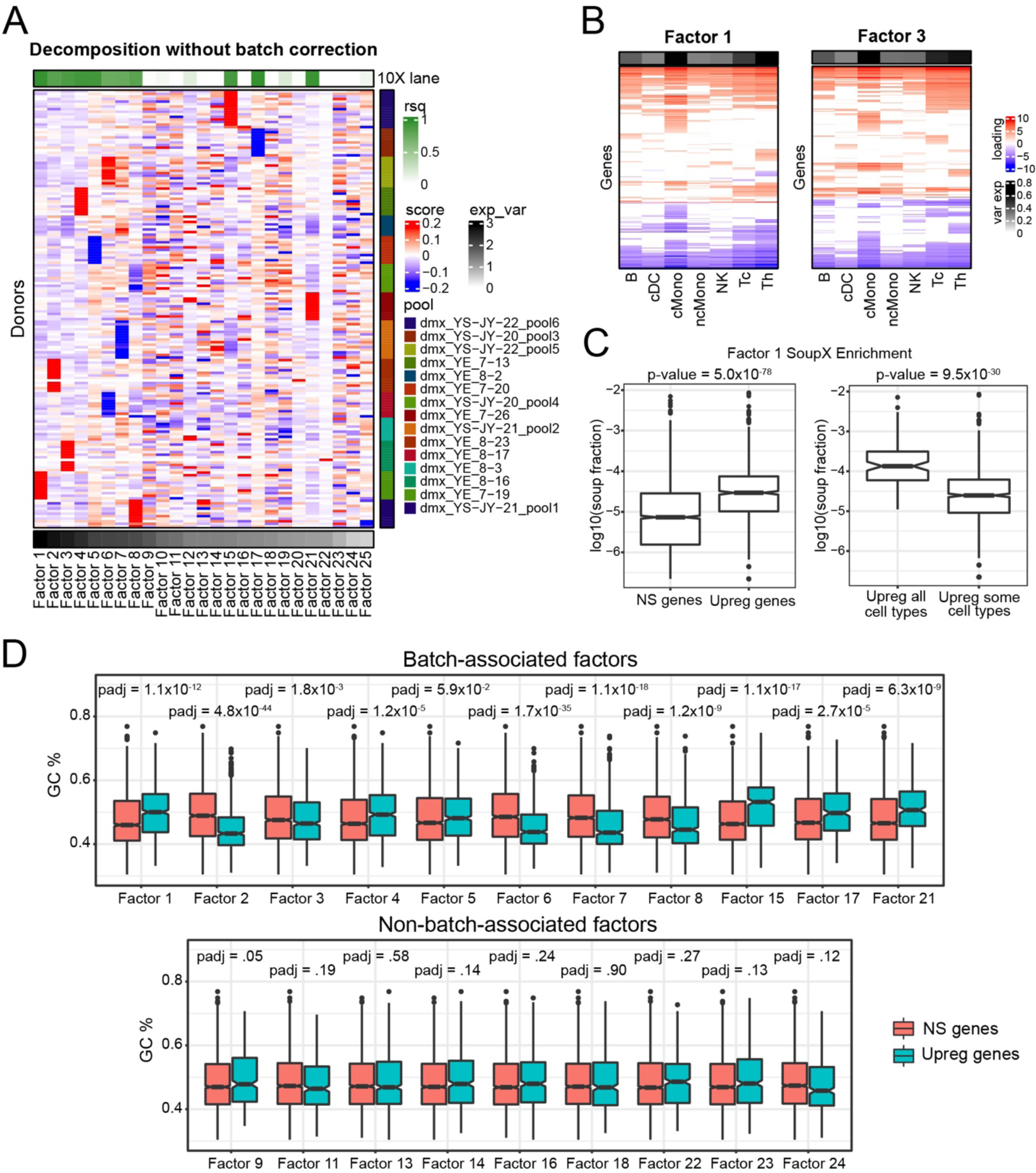
Analysis of technical effects through extracted batch-associated factors. (A) Donor scores matrix for a decomposition of the SLE dataset without applying batch-effect removal. Also shown are p-values for associations between the factor scores and 10X lane (top). The p-values were calculated using univariate linear model F-tests. Columns are ordered by explained variance, shown as a bottom annotation. Rows are grouped by 10X lane, shown as an annotation on the right side. (B) Loadings matrices for two factors associated with 10X Chromium lanes limited to the significant genes only. Rows are hierarchically clustered. (C) Analysis of factor 1 loadings for association with ambient RNA content of batch dmx_YE_7-19. The left plot shows the association between all genes that are upregulated in this batch and their fractional representation in the ambient RNA “soup” for the batch. The right plot shows ambient RNA fractions for genes that were significantly upregulated across all cell types compared to those upregulated only in some cell types. Associations are calculated by two-sample t-tests for (G-H). “NS genes” for (G-H) includes all genes that were not significantly upregulated. (D) Associations between upregulated genes in each factor and GC content. This is shown separately for the batch-associated factors (top) or non-batch-associated factors (bottom).

## Notes

### Summary of Updates

We have included additional validation analyses, supporting the robustness of our findings in SLE. We also included new comparisons between SLE and COVID-19, highlighting conserved and disease-specific processes identified by scITD. Lastly, we included a comparison of our ligand-receptor inference strategy to that of an existing method.

## References

1. Consortium, T.Gte. (2015). The Genotype-Tissue Expression (GTEx) pilot analysis: Multitissue gene regulation in humans. Science 348, 648–660. 10.1126/science.1262110.

2. Melé, M., Ferreira, P.G., Reverter, F., DeLuca, D.S., Monlong, J., Sammeth, M., Young, T.R., Goldmann, J.M., Pervouchine, D.D., Sullivan, T.J., et al. (2015). The human transcriptome across tissues and individuals. Science 348, 660–665. 10.1126/science.aaa0355.

3. Kang, H.M., Subramaniam, M., Targ, S., Nguyen, M., Maliskova, L., McCarthy, E., Wan, E., Wong, S., Byrnes, L., Lanata, C.M., et al. (2018). Multiplexed droplet single-cell RNA-sequencing using natural genetic variation. Nature Biotechnology 36, 89–94. 10.1038/nbt.4042.

4. Stoeckius, M., Zheng, S., Houck-Loomis, B., Hao, S., Yeung, B.Z., Mauck, W.M., Smibert, P., and Satija, R. (2018). Cell Hashing with barcoded antibodies enables multiplexing and doublet detection for single cell genomics. Genome Biol 19, 224. 10.1186/s13059-018-1603-1.

5. Butler, A., Hoffman, P., Smibert, P., Papalexi, E., and Satija, R. (2018). Integrating single-cell transcriptomic data across different conditions, technologies, and species. Nat Biotechnol 36, 411–420. 10.1038/nbt.4096.

6. Haghverdi, L., Lun, A.T.L., Morgan, M.D., and Marioni, J.C. (2018). Batch effects in single-cell RNA-sequencing data are corrected by matching mutual nearest neighbors. Nat Biotechnol 36, 421–427. 10.1038/nbt.4091.

7. Barkas, N., Petukhov, V., Nikolaeva, D., Lozinsky, Y., Demharter, S., Khodosevich, K., and Kharchenko, P.V. (2019). Joint analysis of heterogeneous single-cell RNA-seq dataset collections. Nature Methods 16, 695–698. 10.1038/s41592-019-0466-z.

8. Stuart, T., Butler, A., Hoffman, P., Hafemeister, C., Papalexi, E., Mauck, W.M., Hao, Y., Stoeckius, M., Smibert, P., and Satija, R. (2019). Comprehensive Integration of Single-Cell Data. Cell 177, 1888–1902.e21. 10.1016/j.cell.2019.05.031.

9. Chen, S., Rivaud, P., Park, J.H., Tsou, T., Charles, E., Haliburton, J.R., Pichiorri, F., and Thomson, M. (2020). Dissecting heterogeneous cell populations across drug and disease conditions with PopAlign. PNAS 117, 28784–28794.

10. Crowell, H.L., Soneson, C., Germain, P.-L., Calini, D., Collin, L., Raposo, C., Malhotra, D., and Robinson, M.D. (2020). muscat detects subpopulation-specific state transitions from multi-sample multi-condition single-cell transcriptomics data. Nature Communications 11, 6077. 10.1038/s41467-020-19894-4.

11. Squair, J.W., Gautier, M., Kathe, C., Anderson, M.A., James, N.D., Hutson, T.H., Hudelle, R., Qaiser, T., Matson, K.J.E., Barraud, Q., et al. (2021). Confronting false discoveries in single-cell differential expression. Nat Commun 12, 5692. 10.1038/s41467-021-25960-2.

12. Mathys, H., Davila-Velderrain, J., Peng, Z., Gao, F., Mohammadi, S., Young, J.Z., Menon, M., He, L., Abdurrob, F., Jiang, X., et al. (2019). Single-cell transcriptomic analysis of Alzheimer’s disease. Nature 570, 332–337. 10.1038/s41586-019-1195-2.

13. Corridoni, D., Antanaviciute, A., Gupta, T., Fawkner-Corbett, D., Aulicino, A., Jagielowicz, M., Parikh, K., Repapi, E., Taylor, S., Ishikawa, D., et al. (2020). Single-cell atlas of colonic CD8 + T cells in ulcerative colitis. Nature Medicine 26, 1480–1490. 10.1038/s41591-020-1003-4.

14. Liu, C., Martins, A.J., Lau, W.W., Rachmaninoff, N., Chen, J., Imberti, L., Mostaghimi, D., Fink, D.L., Burbelo, P.D., Dobbs, K., et al. (2021). Time-resolved systems immunology reveals a late juncture linked to fatal COVID-19. Cell 184, 1836–1857.e22. 10.1016/j.cell.2021.02.018.

15. Ren, X., Wen, W., Fan, X., Hou, W., Su, B., Cai, P., Li, J., Liu, Y., Tang, F., Zhang, F., et al. (2021). COVID-19 immune features revealed by a large-scale single-cell transcriptome atlas. Cell 184, 1895–1913.e19. 10.1016/j.cell.2021.01.053.

16. van der Wijst, M.G.P., Vazquez, S.E., Hartoularos, G.C., Bastard, P., Grant, T., Bueno, R., Lee, D.S., Greenland, J.R., Sun, Y., Perez, R., et al. (2021). Type I interferon autoantibodies are associated with systemic immune alterations in patients with COVID-19. Science Translational Medicine 13, eabh2624. 10.1126/scitranslmed.abh2624.

17. Fava, A., and Petri, M. (2019). Systemic Lupus Erythematosus: Diagnosis and Clinical Management. J Autoimmun 96, 1–13. 10.1016/j.jaut.2018.11.001.

18. Allen, M.E., Rus, V., and Szeto, G.L. (2021). Leveraging Heterogeneity in Systemic Lupus Erythematosus for New Therapies. Trends in Molecular Medicine 27, 152–171. 10.1016/j.molmed.2020.09.009.

19. Tucker, L.R. (1966). Some mathematical notes on three-mode factor analysis. Psychometrika 31, 279–311. 10.1007/BF02289464.

20. Langfelder, P., and Horvath, S. (2008). WGCNA: an R package for weighted correlation network analysis. BMC Bioinformatics 9, 559. 10.1186/1471-2105-9-559.

21. Benjamini, Y., and Hochberg, Y. (1995). Controlling the False Discovery Rate: A Practical and Powerful Approach to Multiple Testing. Journal of the Royal Statistical Society: Series B (Methodological) 57, 289–300. 10.1111/j.2517-6161.1995.tb02031.x.

22. Perez, R.K., Gordon, M.G., Subramaniam, M., Kim, M.C., Hartoularos, G.C., Targ, S., Sun, Y., Ogorodnikov, A., Bueno, R., Lu, A., et al. (2022). Single-cell RNA-seq reveals cell type–specific molecular and genetic associations to lupus. Science 376, eabf1970. 10.1126/science.abf1970.

23. Hooks, J.J., Moutsopoulos, H.M., Geis, S.A., Stahl, N.I., Decker, J.L., and Notkins, A.L. (1979). Immune Interferon in the Circulation of Patients with Autoimmune Disease. New England Journal of Medicine 301, 5–8. 10.1056/NEJM197907053010102.

24. Bennett, L., Palucka, A.K., Arce, E., Cantrell, V., Borvak, J., Banchereau, J., and Pascual, V. (2003). Interferon and Granulopoiesis Signatures in Systemic Lupus Erythematosus Blood. Journal of Experimental Medicine 197, 711–723. 10.1084/jem.20021553.

25. Kirou, K.A., Lee, C., George, S., Louca, K., Peterson, M.G.E., and Crow, M.K. (2005). Activation of the interferon-alpha pathway identifies a subgroup of systemic lupus erythematosus patients with distinct serologic features and active disease. Arthritis Rheum 52, 1491–1503. 10.1002/art.21031.

26. Nikpour, M., Dempsey, A.A., Urowitz, M.B., Gladman, D.D., and Barnes, D.A. (2008). Association of a gene expression profile from whole blood with disease activity in systemic lupus erythaematosus. Ann Rheum Dis 67, 1069–1075. 10.1136/ard.2007.074765.

27. Weckerle, C.E., Franek, B.S., Kelly, J.A., Kumabe, M., Mikolaitis, R.A., Green, S.L., Utset, T.O., Jolly, M., James, J.A., Harley, J.B., et al. (2011). Network Analysis of Associations between Serum Interferon Alpha Activity, Autoantibodies, and Clinical Features in Systemic Lupus Erythematosus. Arthritis Rheum 63, 1044–1053. 10.1002/art.30187.

28. Baechler, E.C., Batliwalla, F.M., Karypis, G., Gaffney, P.M., Ortmann, W.A., Espe, K.J., Shark, K.B., Grande, W.J., Hughes, K.M., Kapur, V., et al. (2003). Interferon-inducible gene expression signature in peripheral blood cells of patients with severe lupus. PNAS 100, 2610– 2615. 10.1073/pnas.0337679100.

29. Crow, M.K., Kirou, K.A., and Wohlgemuth, J. (2003). Microarray Analysis of Interferon-regulated Genes in SLE. Autoimmunity 36, 481–490. 10.1080/08916930310001625952.

30. Nehar-Belaid, D., Hong, S., Marches, R., Chen, G., Bolisetty, M., Baisch, J., Walters, L., Punaro, M., Rossi, R.J., Chung, C.-H., et al. (2020). Mapping systemic lupus erythematosus heterogeneity at the single-cell level. Nature Immunology 21, 1094–1106. 10.1038/s41590-020-0743-0.

31. Catalina, M.D., Bachali, P., Geraci, N.S., Grammer, A.C., and Lipsky, P.E. (2019). Gene expression analysis delineates the potential roles of multiple interferons in systemic lupus erythematosus. Commun Biol 2, 1–13. 10.1038/s42003-019-0382-x.

32. Enocsson, H., Wetterö, J., Eloranta, M.-L., Gullstrand, B., Svanberg, C., Larsson, M., Bengtsson, A.A., Rönnblom, L., and Sjöwall, C. (2021). Comparison of Surrogate Markers of the Type I Interferon Response and Their Ability to Mirror Disease Activity in Systemic Lupus Erythematosus. Front Immunol 12, 688753. 10.3389/fimmu.2021.688753.

33. Juárez-Vicuña, Y., Pérez-Ramos, J., Adalid-Peralta, L., Sánchez, F., Martínez-Martínez, L.A., Ortiz-Segura, M. del C., Pichardo-Ontiveros, E., Hernández-Díazcouder, A., Amezcua-Guerra, L.M., Ramírez-Bello, J., et al. (2021). Interferon Lambda 3/4 (IFNλ3/4) rs12979860 Polymorphisms Is Not Associated With Susceptibility to Systemic Lupus Erythematosus, Although It Regulates OASL Expression in Patients With SLE. Frontiers in Genetics 12, 785. 10.3389/fgene.2021.647487.

34. Suen, J.-L., and Chiang, B.-L. (2012). CD4+FoxP3+ regulatory T-cells in human systemic lupus erythematosus. Journal of the Formosan Medical Association 111, 465–470. 10.1016/j.jfma.2012.05.013.

35. Ferreira, R.C., Castro Dopico, X., Oliveira, J.J., Rainbow, D.B., Yang, J.H., Trzupek, D., Todd, S.A., McNeill, M., Steri, M., Orrù, V., et al. (2019). Chronic Immune Activation in Systemic Lupus Erythematosus and the Autoimmune PTPN22 Trp620 Risk Allele Drive the Expansion of FOXP3+ Regulatory T Cells and PD-1 Expression. Front Immunol 10, 2606. 10.3389/fimmu.2019.02606.

36. Casey, K.A., Guo, X., Smith, M.A., Wang, S., Sinibaldi, D., Sanjuan, M.A., Wang, L., Illei, G.G., and White, W.I. (2018). Type I interferon receptor blockade with anifrolumab corrects innate and adaptive immune perturbations of SLE. Lupus Sci Med 5, e000286. 10.1136/lupus-2018-000286.

37. Yung, S., and Chan, T.M. (2015). Mechanisms of Kidney Injury in Lupus Nephritis – the Role of Anti-dsDNA Antibodies. Front Immunol 6, 475. 10.3389/fimmu.2015.00475.

38. Iwata, Y., Wada, T., Furuichi, K., Sakai, N., Matsushima, K., Yokoyama, H., and Kobayashi, K. (2003). p38 Mitogen-Activated Protein Kinase Contributes to Autoimmune Renal Injury in MRL-Faslpr Mice. JASN 14, 57–67. 10.1097/01.ASN.0000037402.83851.5F.

39. Jin, N., Wang, Q., Zhang, X., Jiang, D., Cheng, H., and Zhu, K. (2011). The selective p38 mitogen-activated protein kinase inhibitor, SB203580, improves renal disease in MRL/lpr mouse model of systemic lupus. International Immunopharmacology 11, 1319–1326. 10.1016/j.intimp.2011.04.015.

40. Dodeller, F., and Schulze-Koops, H. (2006). The p38 mitogen-activated protein kinase signaling cascade in CD4 T cells. Arthritis Res Ther 8, 205. 10.1186/ar1905.

41. Wikenheiser, D.J., and Stumhofer, J.S. (2016). ICOS Co-Stimulation: Friend or Foe? Front. Immunol. 0. 10.3389/fimmu.2016.00304.

42. Teichmann, L.L., Cullen, J.L., Kashgarian, M., Dong, C., Craft, J., and Shlomchik, M.J. (2015). Local Triggering of the ICOS Coreceptor by CD11c+ Myeloid Cells Drives Organ Inflammation in Lupus. Immunity 42, 552–565. 10.1016/j.immuni.2015.02.015.

43. Kawamoto, M., Harigai, M., Hara, M., Kawaguchi, Y., Tezuka, K., Tanaka, M., Sugiura, T., Katsumata, Y., Fukasawa, C., Ichida, H., et al. (2006). Expression and function of inducible co-stimulator in patients with systemic lupus erythematosus: possible involvement in excessive interferon-γ and anti-double-stranded DNA antibody production. Arthritis Research & Therapy 8, R62. 10.1186/ar1928.

44. Grimbert, P., Bouguermouh, S., Baba, N., Nakajima, T., Allakhverdi, Z., Braun, D., Saito, H., Rubio, M., Delespesse, G., and Sarfati, M. (2006). Thrombospondin/CD47 Interaction: A Pathway to Generate Regulatory T Cells from Human CD4+CD25− T Cells in Response to Inflammation. The Journal of Immunology 177, 3534–3541. 10.4049/jimmunol.177.6.3534.

45. Tarr, T., Dérfalvi, B., Győri, N., Szántó, A., Siminszky, Z., Malik, A., Szabó, A.J., Szegedi, G., and Zeher, M. (2015). Similarities and differences between pediatric and adult patients with systemic lupus erythematosus. Lupus 24, 796–803. 10.1177/0961203314563817.

46. Stephenson, E., Reynolds, G., Botting, R.A., Calero-Nieto, F.J., Morgan, M.D., Tuong, Z.K., Bach, K., Sungnak, W., Worlock, K.B., Yoshida, M., et al. (2021). Single-cell multi-omics analysis of the immune response in COVID-19. Nat Med 27, 904–916. 10.1038/s41591-021-01329-2.

47. Bastard, P., Rosen, L.B., Zhang, Q., Michailidis, E., Hoffmann, H.-H., Zhang, Y., Dorgham, K., Philippot, Q., Rosain, J., Béziat, V., et al. (2020). Autoantibodies against type I IFNs in patients with life-threatening COVID-19. Science 370, eabd4585. 10.1126/science.abd4585.

48. Buang, N., Tapeng, L., Gray, V., Sardini, A., Whilding, C., Lightstone, L., Cairns, T.D., Pickering, M.C., Behmoaras, J., Ling, G.S., et al. (2021). Type I interferons affect the metabolic fitness of CD8+ T cells from patients with systemic lupus erythematosus. Nat Commun 12, 1980. 10.1038/s41467-021-22312-y.

49. Cruikshank, W.W., Berman, J.S., Theodore, A.C., Bernardo, J., and Center, D.M. (1987). Lymphokine activation of T4+ T lymphocytes and monocytes. The Journal of Immunology 138, 3817–3823.

50. Winkler, M.S., Rissiek, A., Priefler, M., Schwedhelm, E., Robbe, L., Bauer, A., Zahrte, C., Zoellner, C., Kluge, S., and Nierhaus, A. (2017). Human leucocyte antigen (HLA-DR) gene expression is reduced in sepsis and correlates with impaired TNFα response: A diagnostic tool for immunosuppression? PLoS One 12, e0182427. 10.1371/journal.pone.0182427.

51. Olwal, C.O., Nganyewo, N.N., Tapela, K., Djomkam Zune, A.L., Owoicho, O., Bediako, Y., and Duodu, S. (2021). Parallels in Sepsis and COVID-19 Conditions: Implications for Managing Severe COVID-19. Frontiers in Immunology 12, 91. 10.3389/fimmu.2021.602848.

52. Giamarellos-Bourboulis, E.J., Netea, M.G., Rovina, N., Akinosoglou, K., Antoniadou, A., Antonakos, N., Damoraki, G., Gkavogianni, T., Adami, M.-E., Katsaounou, P., et al. (2020). Complex Immune Dysregulation in COVID-19 Patients with Severe Respiratory Failure. Cell Host & Microbe 27, 992–1000.e3. 10.1016/j.chom.2020.04.009.

53. Spinetti, T., Hirzel, C., Fux, M., Walti, L.N., Schober, P., Stueber, F., Luedi, M.M., and Schefold, J.C. (2020). Reduced Monocytic Human Leukocyte Antigen-DR Expression Indicates Immunosuppression in Critically Ill COVID-19 Patients. Anesth Analg 131, 993–999. 10.1213/ANE.0000000000005044.

54. Robinson, M.D., McCarthy, D.J., and Smyth, G.K. (2010). edgeR: a Bioconductor package for differential expression analysis of digital gene expression data. Bioinformatics 26, 139–140. 10.1093/bioinformatics/btp616.

55. Barkas, N., Petukhov, V., Kharchenko, P.V., and Biederstedt, Evan (2021). pagoda2: Single Cell Analysis and Differential Expression.

56. Li, J., Bien, J., and Wells, M.T. (2018). rTensor: An R Package for Multidimensional Array (Tensor) Unfolding, Multiplication, and Decomposition. Journal of Statistical Software 87, 1–31. 10.18637/jss.v087.i10.

57. Sheehan, B.N., and Saad, Y. (2007). Higher Order Orthogonal Iteration of Tensors (HOOI) and its Relation to PCA and GLRAM. In Proceedings of the 2007 SIAM International Conference on Data Mining Proceedings. (Society for Industrial and Applied Mathematics), pp. 355–365. 10.1137/1.9781611972771.32.

58. Kolda, T., and Bader, B. (2009). Tensor Decompositions and Applications. SIAM Rev. 10.1137/07070111X.

59. Unkel, S., Hannachi, A., Trendafilov, N.T., and Jolliffe, I.T. (2011). Independent Component Analysis for Three-Way Data With an Application From Atmospheric Science. JABES 16, 319–338. 10.1007/s13253-011-0055-9.

60. Zhou, G., and Cichocki, A. (2012). Fast and Unique Tucker Decompositions via Multiway Blind Source Separation. Bulletin of the Polish Academy of Sciences: Technical Sciences 60. 10.2478/v10175-012-0051-4.

61. Zappia, L., Phipson, B., and Oshlack, A. (2017). Splatter: simulation of single-cell RNA sequencing data. Genome Biology 18, 174. 10.1186/s13059-017-1305-0.

62. Wolf, F.A., Angerer, P., and Theis, F.J. (2018). SCANPY: large-scale single-cell gene expression data analysis. Genome Biology 19, 15. 10.1186/s13059-017-1382-0.

63. Johnson, W.E., Li, C., and Rabinovic, A. (2007). Adjusting batch effects in microarray expression data using empirical Bayes methods. Biostatistics 8, 118–127. 10.1093/biostatistics/kxj037.

64. Badea, L. (2008). Extracting gene expression profiles common to colon and pancreatic adenocarcinoma using simultaneous nonnegative matrix factorization. Pac Symp Biocomput, 267–278.

65. Korotkevich, G., Sukhov, V., Budin, N., Shpak, B., Artyomov, M.N., and Sergushichev, A. (2021). Fast gene set enrichment analysis. bioRxiv, 060012. 10.1101/060012.

66. Petukhov, V., Igolkina, A., Rydbirk, R., Mei, S., Christoffersen, L., Khodosevich, K., and Kharchenko, P.V. (2022). Case-control analysis of single-cell RNA-seq studies. 2022.03.15.484475. 10.1101/2022.03.15.484475.

67. Satija, R., Farrell, J.A., Gennert, D., Schier, A.F., and Regev, A. (2015). Spatial reconstruction of single-cell gene expression data. Nat Biotechnol 33, 495–502. 10.1038/nbt.3192.

68. Egozcue, J.J., Pawlowsky-Glahn, V., Mateu-Figueras, G., and Barceló-Vidal, C. (2003). Isometric Logratio Transformations for Compositional Data Analysis. Mathematical Geology 35, 279–300. 10.1023/A:1023818214614.

69. Jin, S., Guerrero-Juarez, C.F., Zhang, L., Chang, I., Ramos, R., Kuan, C.-H., Myung, P., Plikus, M.V., and Nie, Q. (2021). Inference and analysis of cell-cell communication using CellChat. Nature Communications 12, 1088. 10.1038/s41467-021-21246-9.

70. Wang, Y., Hicks, S.C., and Hansen, K.D. (2020). Co-expression analysis is biased by a mean-correlation relationship 10.1101/2020.02.13.944777.

71. Gu, Z., Eils, R., and Schlesner, M. (2016). Complex heatmaps reveal patterns and correlations in multidimensional genomic data. Bioinformatics 32, 2847–2849. 10.1093/bioinformatics/btw313.

